# Topological stratification of continuous genetic variation in large biobanks

**DOI:** 10.1101/2023.07.06.548007

**Authors:** Alex Diaz-Papkovich, Shadi Zabad, Chief Ben-Eghan, Luke Anderson-Trocmé, Georgette Femerling, Vikram Nathan, Jenisha Patel, Simon Gravel

**Affiliations:** Quantitative Life Sciences, McGill University, Montreal; Department of Human Genetics, McGill University, Montreal; School of Computer Science, McGill University, Montreal; Department of Bioengineering, McGill University, Montreal

## Abstract

Biobanks now contain genetic data from millions of individuals. Dimensionality reduction, visualization and clustering are standard when exploring data at these scales; while efficient and tractable methods exist for the first two, clustering remains challenging because of uncertainty about sources of population structure. In practice, clustering is commonly performed by drawing shapes around dimensionally reduced data or assuming populations have a “type” genome. We propose a method of clustering data with topological analysis that is fast, easy to implement, and integrates with existing pipelines. The approach is robust to the presence of sub-populations of varying sizes and wide ranges of population structure patterns. We use UMAP and HDBSCAN, respectively methods of dimensionality reduction and density clustering, on data from three biobanks. We illustrate how topological genetic strata can help us understand structure within biobanks, evaluate distributions of genotypic and phenotypic data, examine polygenic score transferability, identify potential influential alleles, and perform quality control.

## Introduction

Following improvements in genomic technologies, large-scale biobanks have become commonplace. The Global Biobank Meta-analysis Initiative (GBMI), for example, lists 23 biobanks with genetic data and health records from over 2.2 million individuals[1]. The growth in sample sizes has led to increased potential for scientific findings, with thousands of genetic loci implicated with phenotypes in genome-wide association studies (GWAS), and used to predict disease traits via polygenic scores (PGS). Though the growth of biobanks has fuelled discovery, population structure—the phenomenon in which allele frequencies systematically differ between populations—remains a persistent confounder in GWAS and PGS (e.g. [2, 3]). Many methods in population genetics seek to describe and account for population structure, but the complexity of human history and of biobank recruitment strategies preclude simple model-based approaches from effectively capturing the many determinants of observed genetic variation.

As an alternative, dimensional reduction and visualization are commonly used to examine both discrete and continuous aspects of genetic variation (e.g. [4, 5]). Within the framework of exploratory-confirmatory data analysis, visualization of complex data enables pattern-recognition and the generation and testing of hypotheses[6]. Visualization alone, however, cannot be used for analysis, and visualization techniques are often used as a precursor to stratification. For example, principal component analysis (PCA) can be used to visualize data and individuals within a certain area are commonly deemed to share an ancestry label. In recently admixed populations (i.e., populations who derive ancestry from “source” populations who had been in relative isolation), grouping based on inferred admixture proportions is also common, often with the use of a reference panel as a proxy for the source populations. By definition, PCA-based approaches capture only the axes of variation that expalin the most variance in a sample, and may not work to discern populations with no reference panel, or with complex admixture histories, or small sample sizes[7]. Other approaches cluster based on shared identity-by-descent (IBD) segments or recent genetic relatedness (e.g. [8, 9]). These approaches typically capture finer scale population structure, but are analytically and computationally demanding. Self-declared variables like race and ethnicity are also sometimes used for genetic stratification but are imperfect indicators of genetic ancestry and are no longer recommend as proxies for it [10, 11].

Despite the demand, there is not an effective, fast, and tractable method for stratifying biobank data based on patterns of genetic structure. In practice, researchers often manually group participants into discrete ad hoc “clusters” that they perceive in low-dimensional visualizations, which they use as strata in downstream analyses regarding, e.g., heterogeneity in ancestry and allele frequencies[12], environmental exposures[4], or assessing the performance of PGS[2, 13]. There are many drawbacks to such ad hoc approaches. For example, in cosmopolitan cohorts, there are many subgroups with distinct ancestral histories, leading researchers to manually distinguish between a “majority” cluster and an “everybody else” cluster—often to be discarded due to its heterogeneity[14, 15].

We propose a topological data analysis approach as an alternative. Rather than fitting individuals to a pre-defined notion of a population, a topological approach describes the network of neighbourhoods between data points—here, this would be the network of genetic similarity between individuals. It is well-suited to describe collections of points in high-dimensional space with smooth distributions but with no clear centre or “archetype”. We assume that structure in high-dimensional genetic data can be represented topologically, and can be locally approximated and reconstructed in a low-dimensional space. After reconstructing data in the low-dimensional space, we identify dense clusters of data—i.e., the genetic strata. This approach is unsupervised, requiring neither a number of clusters nor a reference panel, and thus fits naturally with population genetic data, which is sparse and contains numerous sub-populations of unknown and varying sizes, often without *a priori* definitions.

We demonstrate the effectiveness of this approach on three biobanks, showing that we can consistently and effectively identify and characterize sources of population structure in each cohort, and relate many key variables to this structure. We simultaneously identify structured groups as small as 100 individuals and as large as 400, 000 within the same cohort in a matter of seconds, and describe environmental, sociodemographic, and phenotype variation across groups. We use stratification to identify populations for which PCA adjustment fails within a biobank (often admixed populations) and populations for which PGS transferability is poor (often, but not always, populations diverged from the training population). Finally, we highlight the role of topological modelling in quality control, a critical aspect of the fast-growing biobank space.

Topological modelling, which describes data in terms of local neighbourhoods in a high-dimensional space, is therefore a powerful alternative to ancestral component modelling for the description of genetic variation in complex cohorts.

## Methods

Our method works on genotype data, represented by a matrix of allele counts for each individual and genetic variant. To reduce computational burden, we can also perform analyses on genotype data previously projected to any number of principal components (PCs). We use uniform manifold approximation and projection (UMAP)[16], a dimensionality reduction method, and Hierarchical Density-Based Spatial Clustering of Applications with Noise (HDBSCAN), a clustering algorithm. HDBSCAN has been used before on population genetic data directly on PC-reduced data [17, 18]. As we will see, this results in large proportions of individuals discarded as noise. The application of UMAP leads to increased point density and facilitates clustering (Table S7), and a recent implementation of HDBSCAN by Malzer and Baum (HDBSCAN(*E*^),[19]) drastically reduces the number of discarded points.

UMAP seeks to preserve high-dimensional neighbours in the low-dimensional space[16]. The algorithm requires three parameter inputs: the target number of dimensions, the number of nearest neighbours (used to define the size of high-dimensional neighbourhoods to approximate), and the minimum distance between points in the low-dimensional space. We have previously explored its use for visualization in 2 and 3 dimensions[4]. In this work, we will use UMAP both for visualizing data and for preprocessing data for clustering. Both tasks require different parameter choices (Figure 1):

1. For visualization, we reducing data to 2 dimensions and use a relatively high minimum distance (03 to 05), to facilitate human perception and understanding.
2. For clustering, we reduce data to 3 or more dimensions and use a very low minimum distance (near or equal to 0) to facilitate algorithmic identification of dense clumps of data.

**Figure 1:**
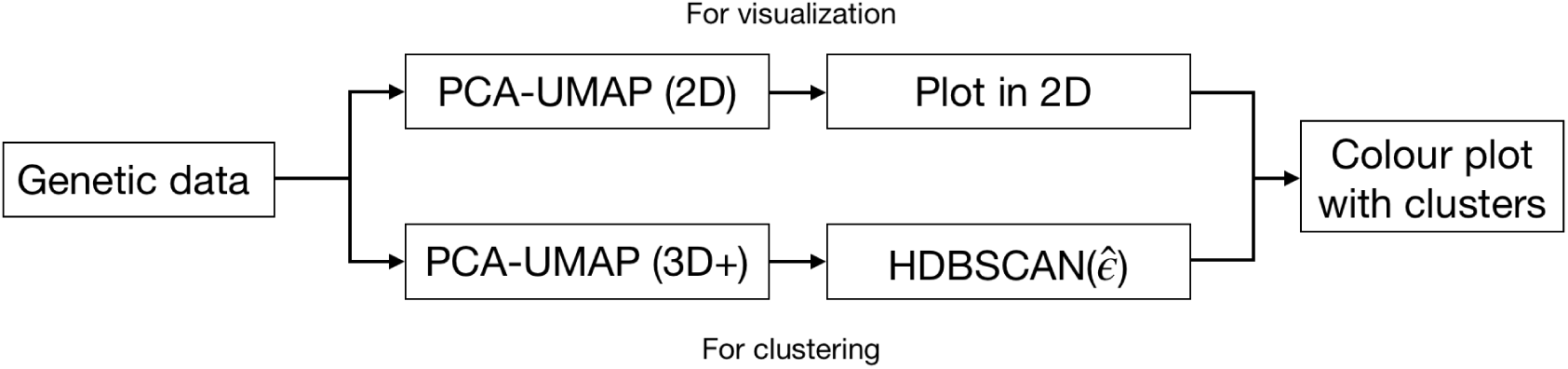
By using different parameters, UMAP can be optimized for clustering or visualization. Visualization requires a two-dimensional representation with the points spread out to make patterns easier to see. In contrast, clustering benefits from UMAP reduction to three or more dimensions to strike a balance between preservation of the original data topology and high point density for clustering. The 2D UMAP plots used to visualize clusterings in this work and are not the same ones used for clustering.

After reducing genetic data to 3 or more dimensions with UMAP in step 2, we use HDBSCAN(*E*^) to extract clusters. HDBSCAN(*E*^) is a hierarchical density-based clustering algorithm based on predecessors HDBSCAN and DBSCAN*[19]. It is motivated by situations where we expect data to be in a sparsely populated space with relatively dense clusters throughout. The number of clusters is not known, and the sizes of the clusters are assumed to vary. This describes biobank data particularly well since they are expected to contain population structure at many different scales, and it is usually difficult to specify in advance a useful number of subgroups to consider. The parameter *E*^ allows clusters to have widely varying sizes; we provide more details on parameters in the Supporting Information (SI).

We use UMAP-assisted density-based clustering on data from three biobanks: the 1000 Genomes Project (1KGP), the UK biobank (UKB), and CARTaGENE (CaG). The 1KGP data consists of the genotypes of 3, 450 individuals sampled from 26 populations from around the world; the populations were decided in advance and their sample sizes are similar, ranging from 104 to 183 samples[20]. The UKB is a cohort of 488, 377 individuals from the United Kingdom (UK) with genotypic, phenotypic, and sociodemographic data. UKB participants were recruited by inviting 9 million individuals registered with the National Health Service (NHS) who lived near a testing centre[21]. CaG is a cohort of residents of the Canadian province of Québec, with genotype data for 29, 337 participants who were recruited using registration data from the Régie de l’assurance maladie du Québec (RAMQ), the provincial health authority, from four metropolitian areas in the province[22]. Unlike the 1KGP, CaG and the UKB do not have *a priori* populations defined, though they collected information about ethnicity, country of birth, and residential geographic distribution.

## Results

### Clustering captures population structure from sample design

The 1KGP’s relatively balanced global sample design makes it useful for testing algorithms to identify population structure. We have previously shown that UMAP results in clear visual clusters from 1KGP data in two dimensions[4]. Figure 2 shows a UMAP representation of the 1KGP. Figure 2a shows the data without population labels (to mimic data with unknown populations), Figure 2b shows the data with corresponding population labels from the 1KGP, and Figure 2c shows the data with cluster labels generated by HDBSCAN(*E*^) run on a 5*D* UMAP.

**Figure 2:**
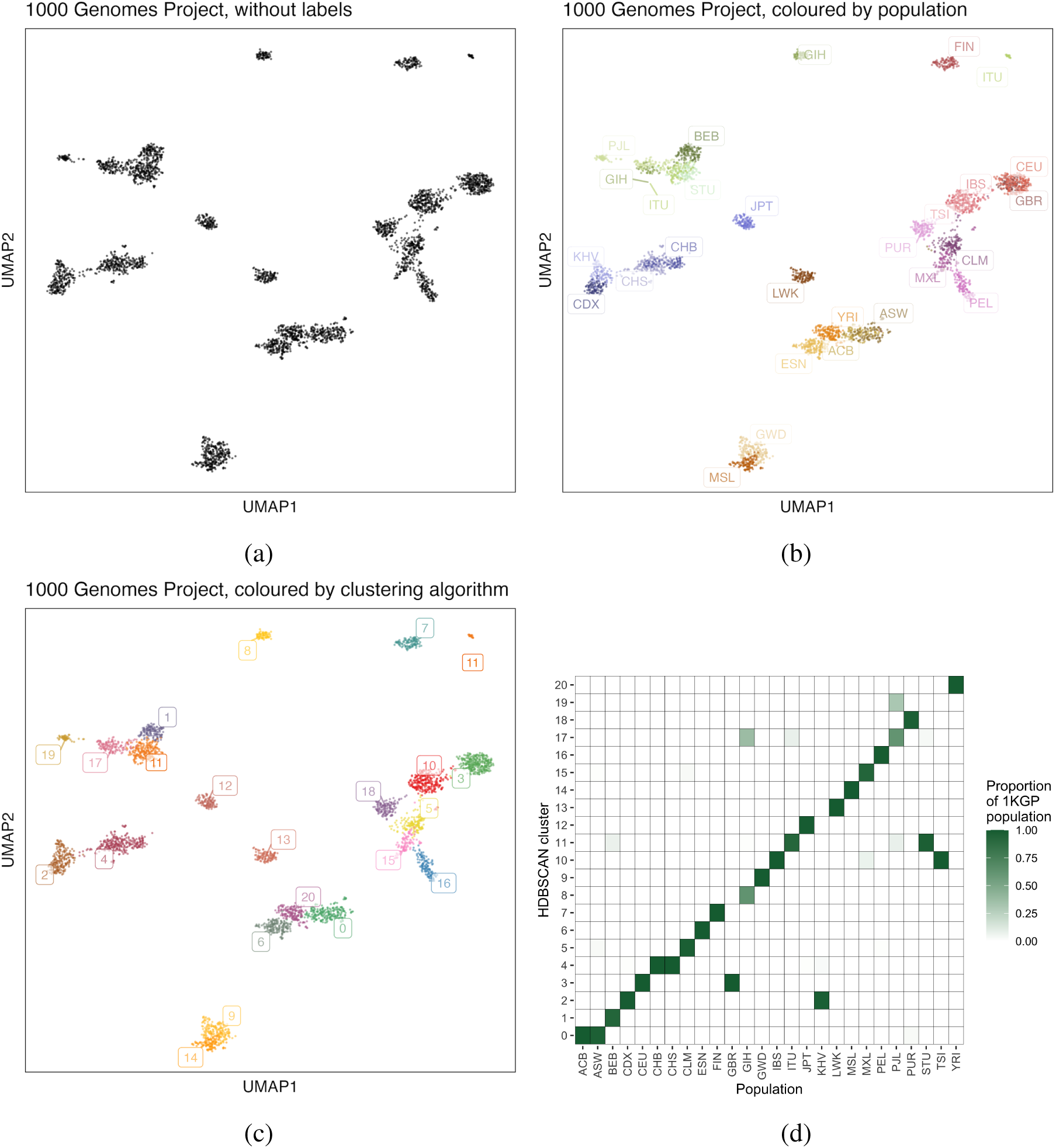
Clusters generated from 1KGP genotype data reflect its population sampling. (A) UMAP embedding of data without labels. (B) UMAP embedding of data, coloured by population label. (C) UMAP embedding of data, coloured by 21 clusters derived from HDBSCAN(*E*^) applied to a 5D UMAP embedding. (D) Proportions of each 1KGP population contained within a given cluster. Most populations fall almost entirely within a single cluster, with a few splitting into multiple clusters. Population labels are provided in Table S4.

The major source of genetic structure in 1KGP data is its sampling scheme, which selected individuals from geographically diverse populations. The clusters formed by UMAP and extracted by HDBSCAN(*E*^) largely reflect this sampling strategy, with some exceptions noted below. Figure 2d shows strong agreement between population label and cluster label, (see also Tables S5 and S6).

These results are comparable to a supervised neural network approach to predict sampled population label (e.g. Figure 3 in [23]), though our approach is unsupervised and runs much more quickly: depending on implementation, deriving the principal components can take 5 to 20 minutes, with the subsequent UMAP step requiring approximately 10 seconds and HDBSCAN(*E*^) less than one second. Comparing these clusters to population labels, the adjusted Rand Index (ARI) is 0769.

**Figure 3:**
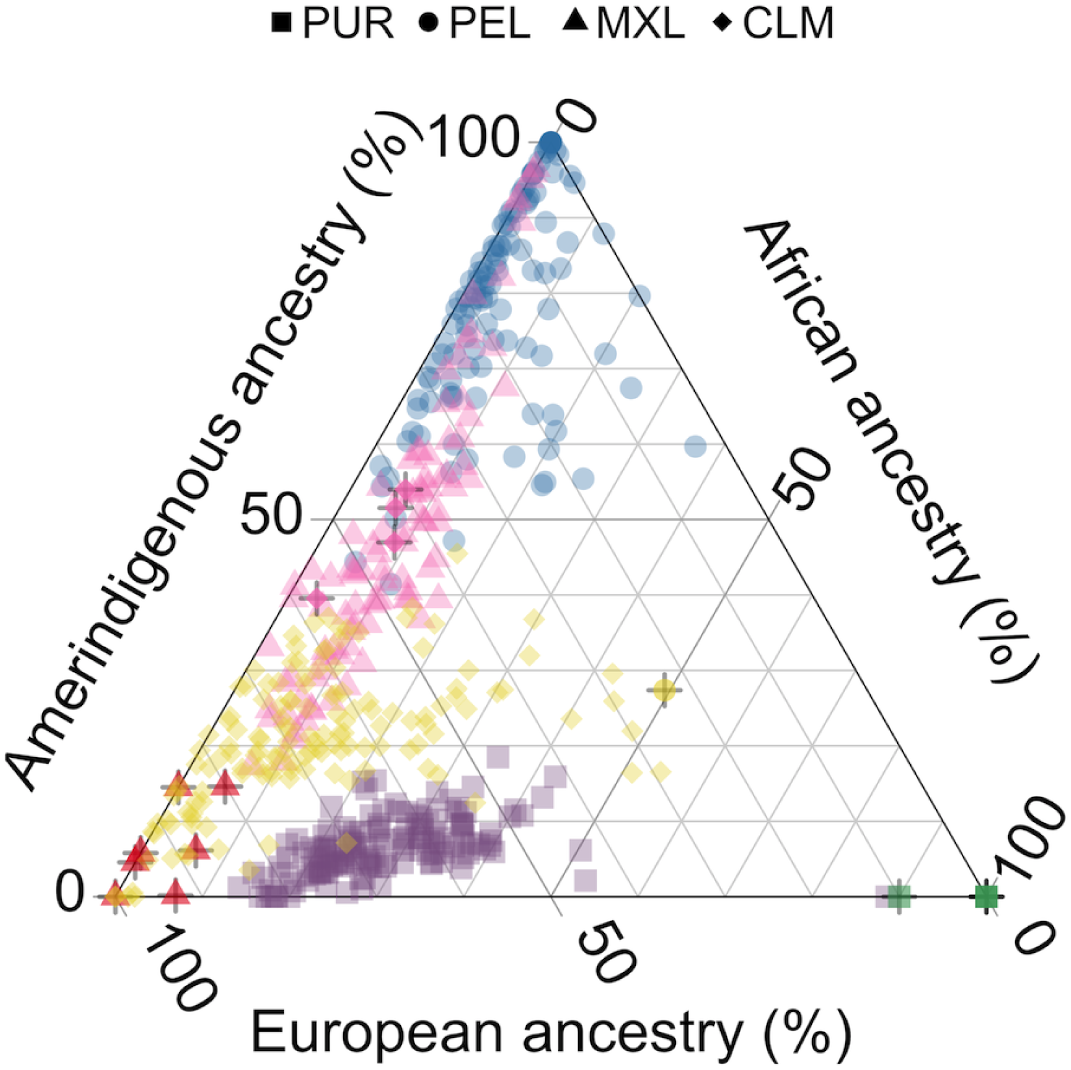
HDBSCAN clusters capture structure in populations with overlapping admixture proportions in the 1KGP. A ternary plot of the PUR, PEL, MXL, and CLM populations from the 1KGP with axes corresponding to global ancestry proportions estimated using ADMIXTURE (*K* = 3). Shapes indicate 1KGP label, colours indicate cluster label and match Figure 2c; bolded points with a + symbol indicate individuals who are not members of the modal cluster of their 1KGP population (full results given in Tables S5 and S6). While many individuals from these populations have similar admixture proportions, UMAP-HDBSCAN(*E*^) is able to extract clusters more effectively than admixture-based methods.

Though there have been methods developed to generate discrete population clusters from genetic data (e.g. [24]), most do not scale to hundreds of thousands of samples. To provide baselines for comparison, we applied k-means clustering to 1KGP data. In one approach we estimated individual-level admixture proportions assuming *K* populations and then applied k-means clustering using the same *K* as the parameter. Using ADMIXTURE with *K* = 21 populations (to match the 21 clusters generated by HDBSCAN(*E*^)), k-means clustering resulted in an ARI of 0611, while *K* = 26 populations (to match the 26 1KGP populations) resulted in an ARI of 0669 (see Figures S1 and S2 for visualizations, and methods for details). In another approach, we applied k-means clustering directly to the leading PCs; the best performing case was *K* = 25 on the top 20 PCs resulting in an ARI of 0696 (Figure S3).

One benefit of the unsupervised approach is that we do not require *a priori* assumptions about the origins of structure, making it possible to capture meaningful clusters despite considerable within-cluster heterogeneity, including in admixed populations. The admixed American population clusters largely match their 1KGP labels (CLM, MXL, PEL, PUR; ARI=0952), despite their heavily overlapping distributions in admixture proportions, illustrated in Figure 3, which is higher than any k-means based clustering (although k-means with k=21 was close at ARI=0.934, see Figure S2b).

Some populations are clustered together: GBR and CEU (British From England and Scotland; and Utah residents with Northern/Western European ancestry), CDX and KHV (Chinese Dai in Xishuangbanna, China; and Kinh in Ho Chi Minh City, Vietnam), IBS and TSI (Iberian Populations in Spain; and Toscani in Italy), ACB and ASW (African Caribbean in Barbados; and African Ancestry in SW USA). While these groups differ in their sampling and history, supervised learning methods also struggle in distinguishing most of these pairs (Figure 3A in [23]). We also note that the CDX and KHV (Cluster 2 in Figure 2b) populations are present at opposite ends of one continuous cloud of points. In other words, two groups belonging to one cluster does not mean that the groups are indistinguishable. Rather, it means that HDBSCAN(*E*^) could find a relatively continuous path in genetic space linking individuals sampled in one group to individuals sampled in the other.

Some South Asian populations are split into different clusters, possibly from stronger patterns of relatedness within those groups[4, 25]. We note the ITU (Indian Telugu in the UK) population is visibly split into two groups in 2D, while clustering carried out in 5D groups them together (Cluster 11). While some clusters will tend to persist across many parametrizations of UMAP and HDBSCAN(*E*^), others based on more subtle patterns or in populations with more continuous variation will be less stable—though discrete groupings can help us understand data, the delineations are always, to a degree, arbitrary.

### Correlates between populations and sociodemographic, phenotypic, and environmental variables

The UK biobank (UKB) contains 488, 377 genotypes from volunteers with an array of demographic, phenotypic, and biomedical data, with individuals’ ages ranging from 40 to 69. The demographic data collected for the UKB include Country of Birth (COB) and Ethnic Background (EB), which is selected from a nested set of pre-determined options (see Table S8). Participants first select their “ethnic group” from a list (e.g. “White”; “Black or Black British”), which determines the list of possible “ethnic background” values (e.g. “British”; “Caribbean”). The most common countries of birth in the data set are England, Scotland, Wales, and the Republic of Ireland, comprising 778%, 80%, 44%, and 10%, respectively. For EB, 883% of participants selected “White British”, with an additional 58% selecting “White Irish” or “Any other white background”. Here we primarily focus on the 28, 814 individuals with other backgrounds.

Many studies of the UKB discard non-European samples, sometimes citing concerns related to confounding from population structure[14]. The population structure has been deeply explored, though typically focused on British or European individuals[26, 27, 28]. Because its sub-populations are numerous, geographically/ancestrally diverse, and of widely varying sizes, clustering the UKB data is challenging, requiring overly broad categorization (e.g. a small number of continental populations [12, 17]) and/or significant computational resources. The original implementation of HDBSCAN, without the *E*^ parameter, discards much of the UKB data as noise and splits populations into hundreds of microclusters that are not interpretable (Fig S4).

Figure 4a shows 26 clusters generated by HDBSCAN(*E*^), placing 9999% of individuals in clusters. We generated word clouds for COB and EB, shown in Figures 4b and 4c, which allow us to illustrate sources of structure without having to impose a label to groups which may be heterogeneous. Individuals in Cluster 10, for example, are mostly born in Somalia (84%), while those in Cluster 23 are mostly born in East Africa (Ethiopia, Sudan, Eritrea; 33%, 29%, 25%, respectively). Those in Cluster 18 are mostly born in sub-Saharan Africa, and 77% chose “African” as their EB, while 19% chose “Other ethnic group”. Figure S5 presents word clouds for another subset of data. Individuals in Cluster 0 are mostly born in Japan and South Korea (84% and 9%, respectively), and those in Cluster 15 are mostly born in Nepal (80%). In contrast, individuals in Cluster 13 are born in a variety of East/Southeast Asian jurisdictions; the most common EB was “Chinese” (70%), followed by “Other ethnic group” (16%) and “Any other Asian background” (11%). Tables S9 and S10 provide breakdowns for clusters.

**Figure 4:**
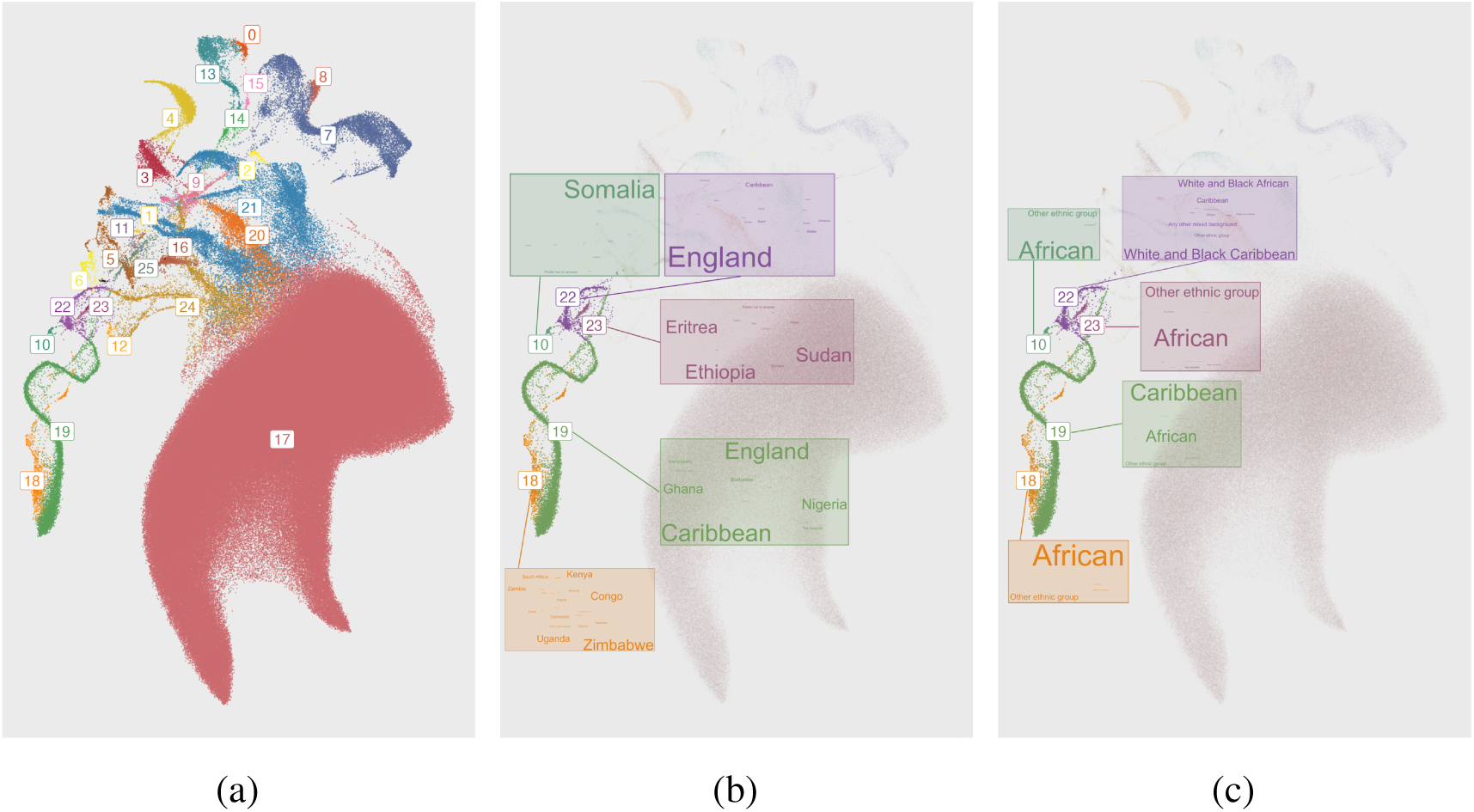
An example of clusters of population structure in the UKB. The clusters reflect a mixture of demographic history within the UK, the geographic origins of recent immigrants, the colonial history of the British Empire, and ongoing admixture. (a) Left: A 2D UMAP of UKB genotypes coloured by HDBSCAN(*E*^). This parametrization generated 26 clusters. (b) Middle: Five clusters are highlighted with word clouds for the most common countries of birth within the cluster. (c) Right: The same five clusters are highlighted with word clouds for the most common EB within the cluster. Admixture proportions for clusters are presented in Figure S6. Detailed breakdowns of EB and country of birth are presented in Tables S9 and S10. An alternative clustering is presented in Figure S15.

Clusters 14 and 22 both capture structure resulting from recent admixture following immigration and colonial history, with 49% and 66% of their respective populations being born in England (see also Figure S6). No single EB represents a majority in either cluster; the most common EB in Cluster 14 is “Any other mixed background” (29%), while for Cluster 22 it is “Mixed, White and Black Caribbean” (39%).

Significant proportions of majority-African-born clusters chose “Other ethnic group” as their EB—a respective 24%, 19%, and 37% in Clusters 10, 18, and 2—suggesting that filtering data based on EB would reduce both genetic and ethnic diversity in a sample. Cluster 18 captures individuals born in sub-Saharan Africa, while Cluster 19 consists of individuals born in the Caribbean (31%), England (28%), as well as Nigeria (14%) and Ghana (12%). These regions are historically linked to the UK; between the years 1641 and 1808, an estimated 325, 311 Africans from the Bight of Benin, between the coasts of modern-day Ghana and Nigeria, were enslaved by British ships and sent to the British Caribbean[29, 30].

Despite the complexity of the UKB, topological clustering identifies population structure that is interpretable from historical or demographic perspectives and includes all or almost all individuals. Such structure is difficult to infer from a single label such as geography or ethnicity; once it is characterized, it can clarify the genetic structure of the cohort.

### Phenotype smoothing and modelling

Epidemiological research often focuses on observed differences between groups—for example, finding the mean of a phenotype or sociodemographic measure and comparing between populations. Clustering is one method to define groups based on shared genetic ancestry, and compare means across groups. However, clustering data featuring continuous variation patterns can be sensitive to input parameters, and the size of clusters can vary across parameterizations, making it challenging to identify the “right” choice of parameters to test for heterogeneity. One approach to visualize heterogeneity across parameter choices is to average phenotypic mean values over multiple parameterization (see Algorithm 1).

To avoid depending on a single parametrization for identifying patterns in phenotypic and sociodemographic data, we can smooth the data over multiple clustering parametrizations (we use 288, outlined in the Supporting Information) using Algorithm 1 and use the smoothed values. This approach has room for improvement but illustrates a proof of concept of incorporating multiple runs of discrete clustering to study patterns in continuous data. Such an approach can be useful in modelling non-linearities in distributions of continuous data and visualize the impact of covariate adjustment in the context of population structure and to identify residual heterogeneity in phenotype distributions (as we present in this section) and environmental data (e.g. smoking rates presented in Figure S8).

**Algorithm 1.**
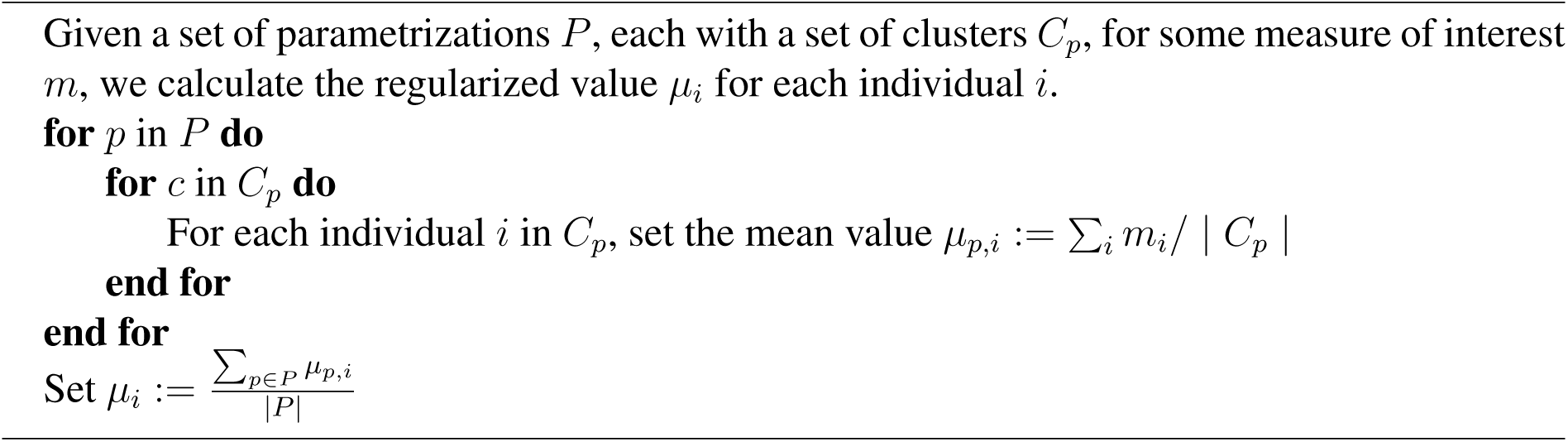
We create a smoothed value for each measure by taking the mean of cluster means for each individual. Given a set of parameters *P* for the clustering algorithm, each parametrization *p* will result in a set of clusters *C_p_*. We use varying cluster assignments across parametrizations to smooth a measured quantity (e.g. phenotype) *m* for individual *i*.

We visualize these smoothed values in Figure 5 for two phenotypes: FEV1 and neutrophil count. Despite having regressed out the effects of the top 40 principal components, there remains structure in the distribution of the residuals, visible at the scale of 05*σ*, where *σ* is the standard deviation of the phenotype across the UKB. For example, the average residual value is noticeably higher in individuals who fell in Cluster 22 as defined in Figure 4a. This cluster is composed mostly of individuals with admixed African/European backgrounds, and although they are intermediate in PCA space to African and European ancestry populations (Figure S11), their phenotype distributions are not intermediate to clusters of primarily European- and African-ancestry individuals (Figure S9, Figure S10).

**Figure 5:**
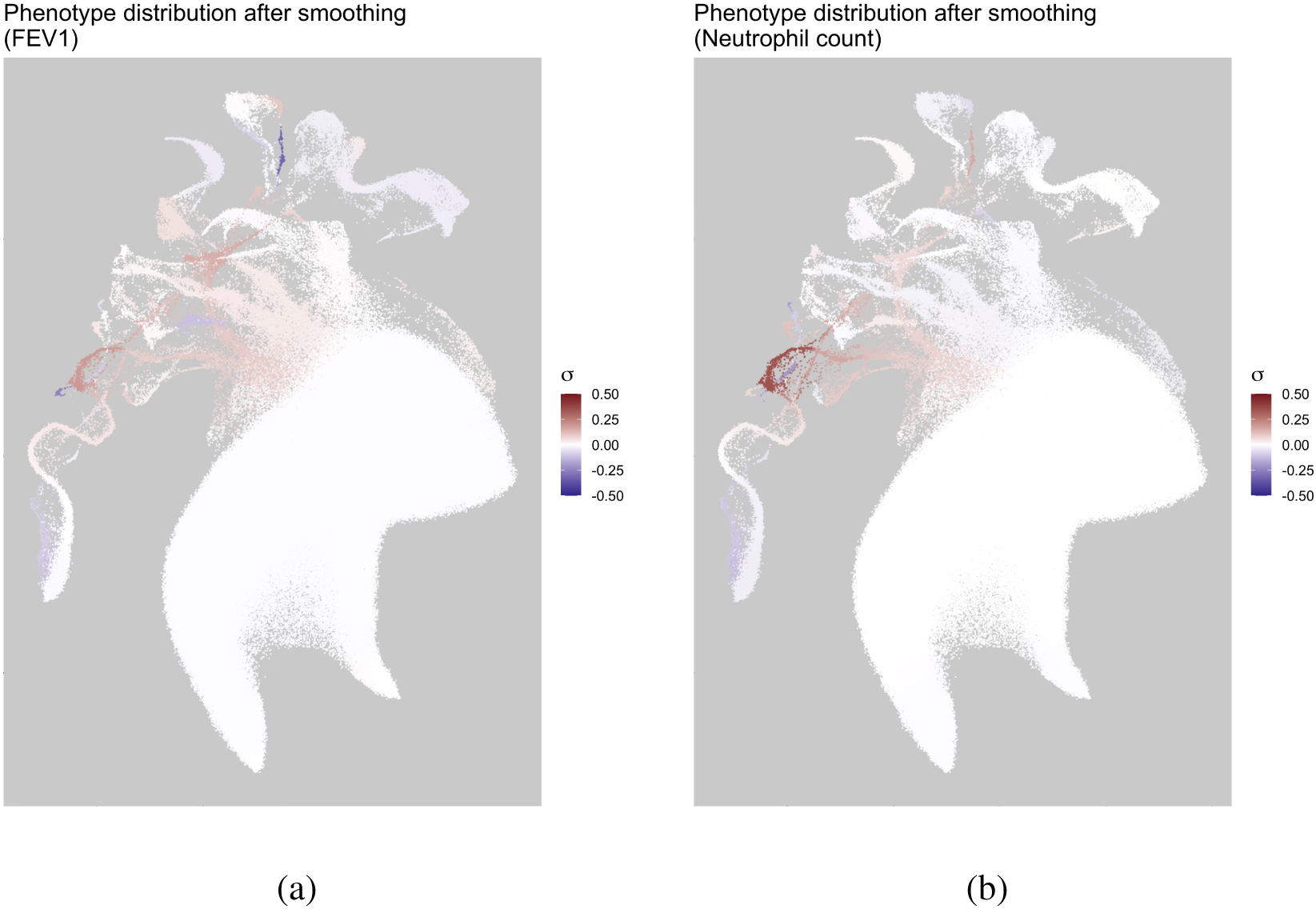
Smoothed phenotypic measures across multiple parametrizations of clustering. A 2D UMAP coloured by phenotype residuals after having regressed the top 40 PCs. Results were averaged by clusters, and we show averaged averages over 288 parametrizations of the clustering pipeline. The colour scale runs from −05*σ* to 05*σ*, for the standard deviation *σ* of each phenotype after regressing the linear effects of the top 40 PCs. We observe that the distributions of phenotypes among some groups are not centred about 0 even after PC adjustment. (a) Left: FEV1. (b) Right: Neutrophil count.

To test if smoothed cluster estimates have explanatory power for these admixed individuals, we compared simple linear models for phenotype prediction using the top 40 PCs versus using the smoothed estimates made from residuals after removing the effects of the top 40 PCs. We compared the models for populations that selected “Mixed” as their EB in the UKB question-naire and found that for individuals who selected “White and Black Caribbean” (*n* = 573) or “White and Black African” (*n* = 389), the smoothed cluster estimates indeed outperformed the PCA model, with an improved mean squared error across several phenotypes (see Figure 6; full table of MSE values in Tables S11 and S12).

**Figure 6:**
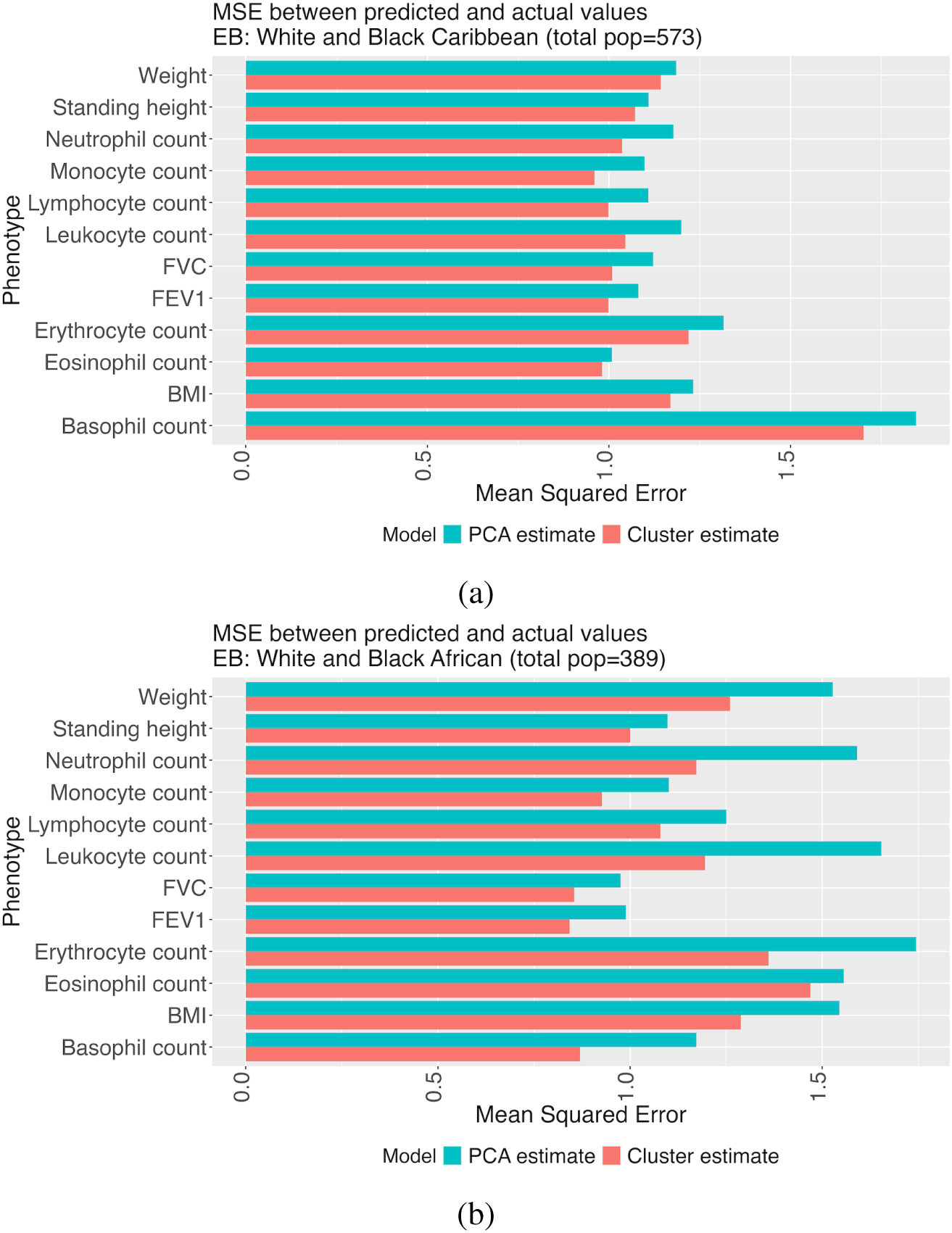
Cluster-based estimation can improve phenotype models. To test the explanatory value of smoothed cluster estimates generated from Algorithm 1, we carried out a five-fold cross-validation on the UKB data, compared phenotype prediction using the top 40 principal components versus estimates generated from the residual structure presented in Figure 5, and calculated the average MSE across folds.

### Evaluating transferability of polygenic scores

Most investigations of PGS transferability are done at a population-level using large-scale geographical groups (e.g. “African”, “European”, “Asian”). However, these broad populations themselves exhibit population structure[31]. To illustrate the value for finer population groupings, we use our 26 cluster labels from Figure 4a, and compared the transferability of PGS across them.

Using UKB data, we estimated effect sizes of SNPs using VIPRS[32]. As a training population, we used individuals who selected “White British” as their EB to mimic the well-documented overrepresentation of European-ancestry individuals in GWAS. We estimated phenotypes for individuals and calculated the values of the fixation index (*F_ST_*) between the clusters. In Figure 7, we plot the PGS accuracy for two phenotypes—standing height and low-density lipoprotein cholesterol (LDL)—against the *F_ST_* for each cluster relative to Cluster 17, a cluster with over 400, 000 individuals and with significant overlap with the training population (*>* 95% selected “White British” as their EB). We observe for height (Figure 7a) that as the *F_ST_* between populations grows, the predictive value of the PGS decreases; such a decrease is expected, due to factors like population-specific causal variants, gene-by-environment interaction, differences in allele frequencies, and linkage disequilibrium between assayed SNPs and causal variants[33].

**Figure 7:**
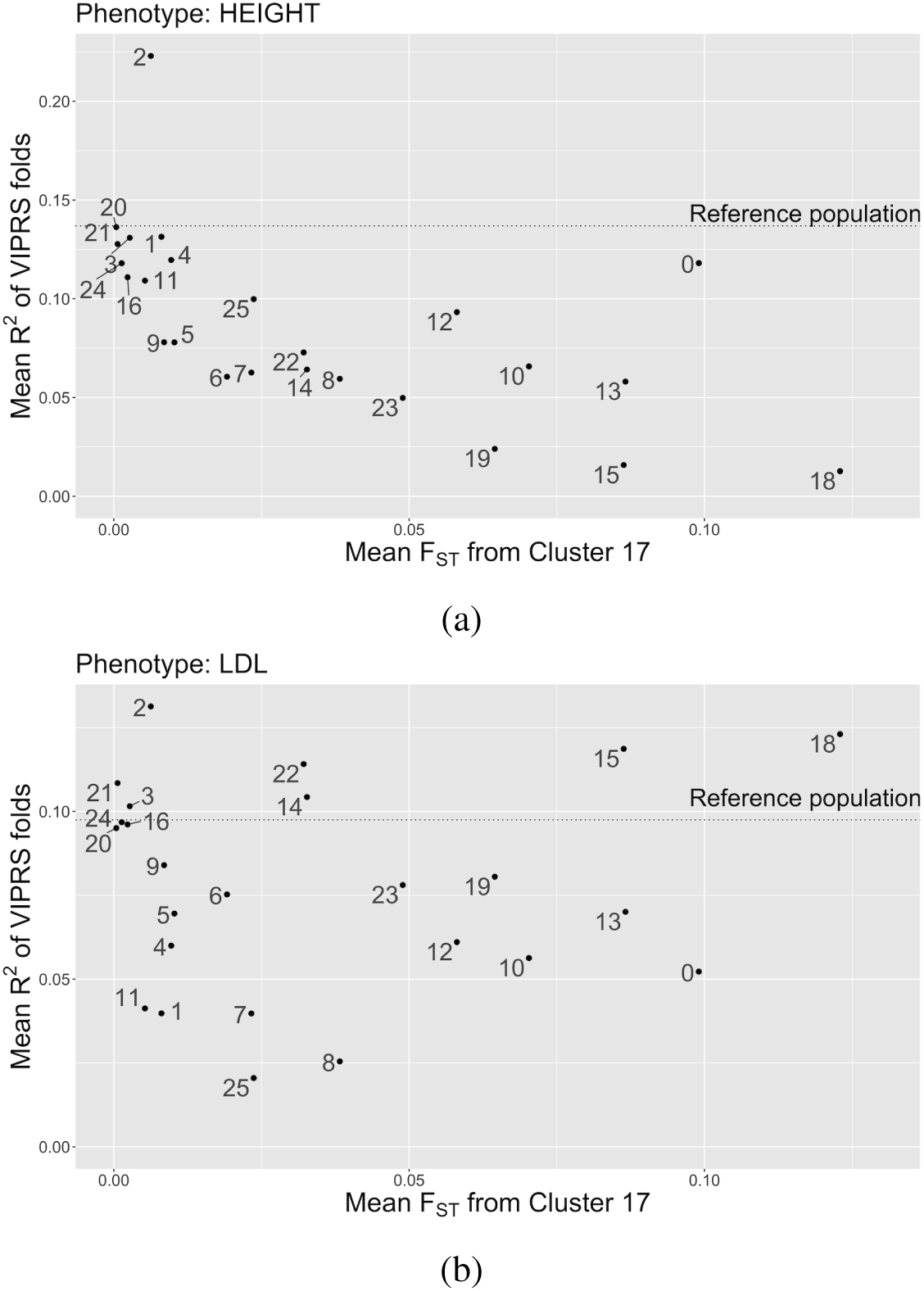
PGS accuracy by *F_ST_* for standing height and LDL. A plot of the mean *R*^2^ of a PGS against the difference in *F_ST_* from the White British in the UKB. We use clusters extracted using HDBSCAN(*E*^). There is a negative linear relationship between *F_ST_* from the largest cluster and PGS accuracy. (a) Top: A PGS of height shows a strong decay between *R*^2^ and *F_ST_*, as expected. (b) Bottom: A PGS of LDL-cholesterol has an unclear relationship between *R*^2^ and *F_ST_* . Cluster 18 has the largest *F_ST_* but one of the highest *R*^2^ values; the cluster also has the highest frequency of the *rs*7412 and *rs*4420638 alleles.

However, we see no such relationship for LDL (Figure 7b). Cluster 18, composed mostly of individuals born in sub-Saharan Africa and of whom 77% selected the EB “Black African”, has one of the best PGS predictions despite its large *F_ST_* from the training population. This may be because there are a few variants with large effect sizes; in contrast to height, LDL has been noted for its relatively low polygenicity[7]. Since *F_ST_*compares genome-wide variation, the accuracy of a PGS constructed from relatively few variants with strong effects is not expected to correlate as strongly with *F_ST_* .

To test if the frequencies of certain alleles impacted the PGS estimates, we modelled the *R*^2^ from the VIPRS estimates for each cluster against minor allele frequencies (MAF) of the top 100 SNPs and found the two strongest results were for *rs*4420638 and *rs*7412 (Tables S1 and S2; Figures S12 and S13, respectively). Both have their highest frequencies in Cluster 18 and both markers are in the apolipoprotein E (APOE) gene cluster; *rs*7412 had the largest overall effect size (*β*^^^ = −01812), while *rs*4420638 had the second largest effect size in the opposite direction (*β*^^^ = 002813). The *rs*7412 allele has been linked to LDL[34] and was found to explain significant variation in LDL in African Americans[35]. The *rs*4420638 allele was associated with LDL even in the presence of the *rs*7412 allele in a study of Sardinian, Norwegian, and Finnish individuals[36]; it was also found to affect LDL in studies of children in Germany[37] and China[38].

The relationship between PGS accuracy and fine-scale population structure is complex and will vary by phenotype. It is not immediately obvious whether a PGS will transfer when there is a large degree of differentiation between the estimand and training populations. However, an approach like UMAP-HDBSCAN(*E*^) can provide a detailed picture of the likely performance of a PGS in various genetic subgroups.

### Quality control for complex multi-ethnic cohorts

Generally the fine-scale structure of biobank data is not known in advance. The structure of under-represented groups in particular, such as minority populations or those with complex histories of recent migration and admixture, can also be intricate and poorly understood, at least by geneticists. Individuals with uncommon combinations of ancestral, geographic, and ethnic descriptors are present in all biobanks. These combinations can be real and represent the completely different nature of genetic ancestry and ethnicity; they may also represent clerical errors[39]. Distinguishing the two is especially relevant when biobanks are used as sample frames for deeper sequencing or for follow-up studies, and when variables like country of birth and ethnicity are used as selection criteria. Using HDBSCAN(*E*^) to explore the relationship between clusters membership and auxiliary variables can detect data collection errors before sample selection is carried out, preventing serious methodology problems or unnecessary exclusion of individuals.

CARTaGENE is a biobank of residents from Quebec, Canada, that has recently genotyped 29, 337 individuals[22]. We were interested in identifying populations of North African descent for further study. In Figure 8, we identified a cluster of 446 people born largely in North Africa with 51 individuals (114%) recorded as being born in American Samoa, an American island territory in the South Pacific Ocean with fewer than 50, 000 inhabitants. After researching possible historical explanations (e.g. migration between American Samoa and North Africa), we traced the result to a coding error from different country codes used over the course of data collection; the actual birth country was corrected to Algeria. The same coding error was found in other clusters, affecting 266 individuals born in 43 countries. While this error was easy to discover using HDBSCAN(*E*^), it is not obvious whether or how it would have been identified otherwise given that it affected less than 1% of the cohort. Efficient data exploration, aided by visualization and clustering, remains one of our best tools to combat the dual evils of bookkeeping errors and batch effects.

**Figure 8:**
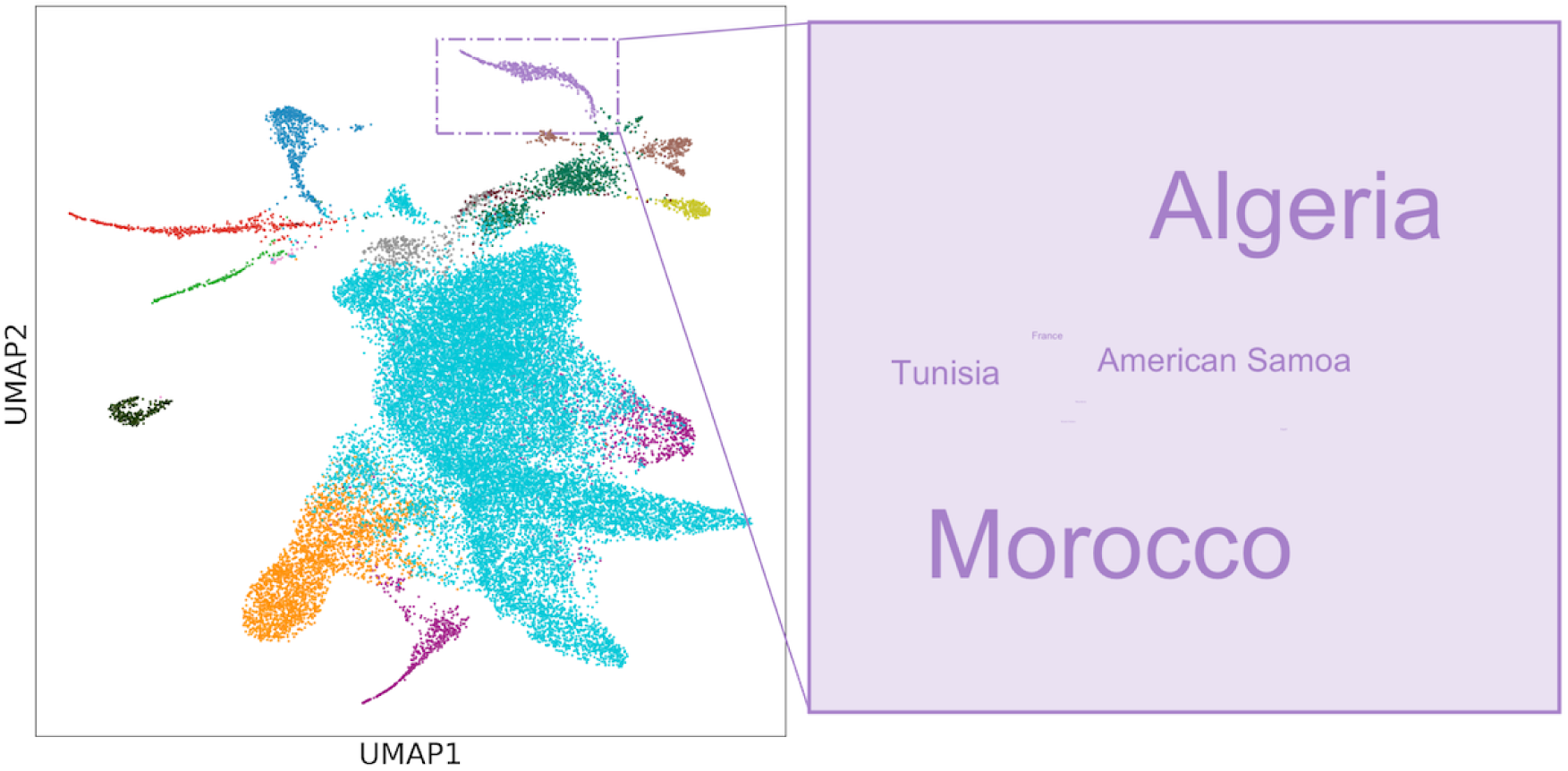
Clustering can identify data collection errors. A 2D UMAP of CARTaGENE data coloured by clusters extracted using HDBSCAN(*E*^). The highlighted cluster was found to have most of its individuals born in North Africa. A word cloud shows that a significant minority of individuals were born in American Samoa, which was found to be a coding error.

## Discussion

We present UMAP-HDBSCAN(*E*^), a new approach to describe population structure that approximates the topology of high-dimensional genetic data and detects dense clusters in a low-dimensional space.

The most commonly used approaches for fine-scale genetic community identification are based on measures of recent relatedness such as identity-by-descent (IBD; see e.g. [8, 9, 40, 41]). An IBD-based approach in ATLAS, for example, recently identified associations between genetic clusters and genetic, clinical, and environmental data [42]. The ability of IBD clustering to identify fine-scale structure can be due to two effects. First, it focuses on recent relatedness between individuals, which may be helpful in identifying recent demographic effects. Second, because it focuses on pairwise similarity, it encourages the use of clustering methods that focus on genetic neighbourhoods, i.e., on more topological approaches.

Despite the existence of such methods, researchers commonly rely on hand-delineation of dimensionally-reduced data (e.g. [12, 2]). This is because IBD clustering is analytically demanding, and because IBD clusters focus on recent relatedness and may not reflect overall genetic similarity observed in PCA or UMAP plots. The topological approach presented here is meant to capture overall genetic similarity. Since it bypasses the need to perform phasing and IBD calling, it requires fewer analytical tools and computational resources: starting from UMAP data, clustering the UKB takes less than 60 seconds on a single core. It can model populations of widely varying sizes and requires neither reference panels nor *a priori* definitions of populations, but can use auxiliary data such as geographic coordinates, jurisdiction, country of birth, population label, ethnicity, etc., to characterize the clusters *a posteriori* and learn about their history or origins.

A recent publication suggested moving entirely away from stratification based on genetic clusters [7]. Instead, they argued in favour of individual-level measures. They cite three issues with clusters: (i) clustering algorithms fail to capture populations without reference panels, such as those that are relatively small or recently admixed; (ii) clusters ignore inter-individual variation; and (iii) clustering results change based on algorithms and reference panels. We believe that these criticisms are valid for the type of archetype-based stratification considered in [7]: if an individual fell within a certain PCA distance of one of nine pre-defined population centroids, they were considered a member of a cluster; otherwise, their ancestry was considered unknown.

We believe that the first two objections can be resolved by topological approaches. In the UKB, 91% of participants were placed into clusters in [7]. In contrast, across 604 runs with varying parameters, the median percentage of individuals placed in a cluster was 9999% (Figure S14), with the three worst-performing runs of UMAP-HDBSCAN(*E*^) respectively assigning 9911%, 9969%, and 9986% of individuals in the UKB to a cluster. The clusters reflect groups that have shared genetic and geographic histories, including for relatively small and recently admixed groups which were often excluded based on prior approaches[7, 15]. We achieved similar results with CaG and 1KGP data, suggesting that our approach is robust to details of biobank composition.

### Applications

Understanding the population structure of a biobank is a necessary precursor to many analyses. In the 1KGP, the source of its structure is largely the sampling scheme, which is reflected in Figure 2—the populations were deliberately sampled from multiple locations around the world with similar sample sizes. The sources of population structure of the UKB, on the other hand, reflect a complex history of migrations at different geographic and time scales, including isolation by distance within the UK and recent immigration and admixture. The structure of a typical biobank is more similar to the UKB than the 1KGP, as the recruitment methodology is often based on residence within a jurisdiction. Examples include municipal (ATLAS in Los Angeles[42], Bio*Me* in New York City[43]), regional (CARTaGENE in Quebec[22]) and national (Million Veterans Project (MVP),[44], CANPATH[45]) biobanks. Leveraging these diverse cohorts can improve variant discovery[46, 47].

Though population labels like ethnicity can be useful, individuals may identify as “Other” or “Unknown”, leading to incomplete data. In the MVP, missing data were imputed using a support vector machine trained on race/ethnicity data to harmonize genetic data with labels for an ethnicity-specific GWAS[48]. A similar supervised approach with random forests was used by gnomAD[18]. Rather than assigning ethnicities to individuals, we constructed clusters from genetic data and investigated the distributions of auxiliary variables within clusters, including missing values. We found word clouds to be well-suited for describing data without imposing a reductive label.

The goal of genetic stratification is in no way to replace self-declared variables in contexts where they are relevant. In fact, genetic stratification revealed interesting trends in self-declared variables. For example, in Cluster 17 of Figure 4a, 976% of individuals were born in Britain and Ireland and 995% chose an ethnic group label; in contrast, 189% of those in Cluster 18 (mostly born in sub-Saharan Africa) and 365% in Cluster 23 (mostly born in the Horn of Africa) chose “Other”, highlighting differential completeness of questionnaire data. UKB strata with “mixed” ethnic backgrounds as their mode featured multiple ethnic background labels, likely reflecting both the fact that (genetically) admixed individuals may have a diversity of ethnic backgrounds, and the fact that individuals with both mixed genetic and cultural heritage may have to choose among potentially inadequate labels (see, e.g., discussion in [15]). The presence or absence of a label in data collection can critically influence how people identify: between the 2011 National Household Survey and the 2016 Census in Canada, there was a 536% drop in people who identified as “Jewish” simply because the label was not provided as an example ethnicity in the 2016 questionnaire[49].

### Considerations

Unlike archetype-based methods, HDBSCAN(*E*^) identifies groups that can be created by linking nearby individuals—it is possible to have a long chain containing many individuals who are each closely related to those near them within a cluster but not to those at the distant end. In this way, admixed populations can form a single cluster even though individuals within cluster can differ as much as individuals from the different ancestral “source” populations. In a sense, HDBSCAN(*E*^) identifies groups of individuals whose distribution in genetic space suggests a common sampling or demographic history, rather than genetic similarity. For this reason, topological stratification may be less conducive to reification of clusters and the notion that population labels reflect a true underlying “type”. However, given the weaponization of population genetics research in the past[50], it is worth emphasizing limitations common to all clustering approaches.

No single label is an individual’s “true” ancestry, race, or ethnicity, as these are complex, multifactorial population descriptors[15, 51]. Thus clustering does not have a well-defined ground truth [52], and clusters are most useful as “helpful constructs that support clarification”[53]. With real genetic data, there is no “correct” number of populations[54] and discrete groupings provide a flattened view of a high-dimensional landscape[15, 55]. The clusters generated are sensitive to the input samples, since the demographic composition of a biobank will impact the clustering, and they are also affected by the parameters at the filtering, dimensionality reduction, and clustering steps. This is a reflection of the data, as genetic data are not composed of “natural types”. These clusters can be useful in understanding how genetics relates to health and the environment, but variation in phenotypes across genetic clusters does not imply a genetic cause, as differences in environment or systemic discrimination are also expected to produce such variation[56]. Each identified cluster is also heterogeneous. The UK biobank clusters of majority sub-Saharan-born individuals, for example, encompass considerable genetic substructure[57]. Different choices of metrics for clustering (i.e., genetic relatedness vs. IBD) can emphasize different types of structure. There are no true clusters.

Ultimately, however, many useful analyses require some definition of “populations”. For example, an allele frequency can only be calculated and reported within a population. Data exploration and quality control often require investigating relevant subsets of the data to decide whether they reflect technical artefacts or meaningful subgroups. To date there has not been a method of stratification that is tractable, easy to implement, robust to the presence of many populations of many sizes, and that captures all or almost all individuals with complex population histories. We believe our topological approach satisfies these important needs. Looking forward, we expect that topological approaches underlying UMAP and HDBSCAN(*E*^) also present a promising avenue to move towards a more continuous description of genetic variation in complex cohorts.

## Acknowledgements

We are grateful to the participants in each biobank who provided their genetic data. We thank the CARTaGENE team for troubleshooting data with us, and C. Bhérer, M. L. Spear, and P. Verdu for scientific discussion.

## Materials and Methods

Our code is available at https://github.com/diazale/topstrat. We have provided command line tools to run Python implementations of UMAP and HDBSCAN(*E*^). We used three datasets for this analysis: the 1000 Genomes project (1KGP), the UK biobank (UKB), and CARTaGENE (CaG).

For the 1KGP we used 3, 450 genotypes using Affy 6.0 genotyping[20]. We generated the principal components using a Python script and have made the top PCs available in the repository to demonstrate the code. We used the genotype file

ALL.wgs.nhgri_coriell_affy_6.20140825.genotypes_has_ped.vcf.gz

and population labels

affy_samples.20141118.panel 20131219.populations.tsv,

available at http://ftp.1000genomes.ebi.ac.uk/vol1/ftp/release/20130502/ supporting/hd_genotype_chip/. We generated admixture proportions using ADMIXTURE 1.3.0[58] from 45, 197 SNPs. Using 32GB of RAM and 32 cores, this took 10, 554 seconds to run with *K* = 21 populations and 40, 719 seconds to run with *K* = 26 populations. For the UKB, we limited our analyses to the 488, 377 individuals with genotype data. We used the UKB’s top 40 pre-computed PCs (Data-Field 22009), blood cell counts (Data-Fields 30000, 30010, 30120, 30130, 30140, 30150, 30160), lung function measures (Data-Fields 3062, 3063), age (Data-Field 21003), sex (Data-Field 31), standing height (Data-Field 50), weight (Data-Field 21002), BMI (Data-Field 21001), smoking status (Data-Field 20116), country of birth (Data-Fields 1647, 20115), and ethnic group/background (Data-Field 21000). Ethnic group/background is a hierarchical item in which participants are prompted to select from a pre-populated list of options for Ethnic Group (e.g. “White”) and, if available, a secondary option for Ethnic Background (e.g. “British”). Phenotypes used in analyses were normalized with respect to variables *sex*, *age*, and *age*^2^. Access to the UKB can be granted at https://www.ukbiobank.ac.uk/scientists-3/genetic-data/.

For CARTaGENE, we used 29, 337 individuals with genotype data. We generated the PCs using PLINK[59] after filtering for linkage disequilibrium and HLA (chromosome 6, 25000000**–**33500000). The options used were:

- indep-pairwise 1000 50 0.1 (PLINK2)
- maf 0.05
- mind 0.1
- geno 0.1
- hwe 1e-6.

We used the Python implementations of UMAP[16] (0.3.6) and HDBSCAN (0.8.24), integrating the updates from Malzer and Baum[19]. To calculate PGS, we used VIPRS[32].

## Supporting Information (SI)

For visualization, we reduce our data to 2D via UMAP and set a relatively high minimum distance (*MD*; usually between 0.3 and 0.5); this enables us to view fine-scale patterns of structure. We find satisfactory results with the number of neighbours (*NN*) varying from 15 to 50; higher values will require more computational resources, but they increase the connectivity between points in the data, as discussed in [60]. For clustering, we set a low value of minimum distance (equal to or close to 0) and reduce the number of dimensions to at least 3—in our analyses, we used 3, 4, and 5 dimensions. The low minimum distance encourages dimensionally-reduced data to form dense clusters, while keeping the dimensionality at ≥ 3 preserves the complexities of data that can be lost because of artificial tearing in the drop from 3 to 2 dimensions. The number of neighbours will vary depending on what is a reasonable expectation for the data. For the 1KGP data, which consists of geographically diverse samples of roughly similar size, 50 neighbours capture the structure well. For biobank data, it is common for structure to arise from a handful of individuals; we found 10 to 25 neighbours to work best. Lower neighbourhood values (e.g. *NN* = 5) will create smaller clusters, but can also highlight highly-localized structure within larger populations. 2*D* visualizations can give intuition as to the presence and sizes of clusters. If pre-processing the data with PCA, more PCs tend to reveal finer-scale structure (see e.g. the relationship with geographical coordinates in Figures S17 and S18 in [4]). For the 1KGP clusters in Figure 2 we used the top 16 PCs; for the UKB in Figure 4a and CaG in Figure 8 we used the top 25.

In parametrizing HDBSCAN(*E*^), the parameter *E*^ defines a threshold at which clusters are merged or split. We find values of *E*^ ranging from 0.3 to 0.5 to be effective at ensuring all or almost all individuals are clustered while still identifying fine-scale structure. The minimum number of points (*MP*) should not be significantly higher than the number of neighbours used in the associated UMAP. If *MP* is high and *NN* is low, it can result in a large number of points being classified as noise since the UMAP data will tend to form small clusters; e.g. a UMAP parametrized with *NN* = 10 and HDBSCAN(*E*^) with *MP* = 100 may return poor results.

Changing parameters will result in different clusters being generated. Given the low computational costs of UMAP and HDBSCAN(*E*^), we recommend running a grid search for visualization and exploratory analysis. Clusters can then be characterized using auxiliary data, such as country of birth, geographical location, population label, self-identification, etc. We selected the clustering for the UKB for its suitability for comparing PGS results by *F_ST_* from the training population. For CaG, we selected one of the clustering runs that generated a cluster of individuals with North African ancestry.

We calculated pairwise *F_ST_* for UKB clusters using PLINK[59]. We calculated admixture proportions using ADMIXTURE 1.3.0[58]. For computational reasons, for the UKB we calculated admixture proportions on individuals not falling into Cluster 17 (the largest cluster, containing around 400, 000 individuals) in Figure 4a.

Visualizations and statistical analyses were done in R (3.5.3)[61]. We used ggplot2[62] for graphics and ggwordcloud for word clouds, and stargazer[63] to generate tables.

For phenotype smoothing, we removed the effects of the top 40 PCs using linear regression, working with the residuals. For phenotype *p* and individuals *i* = 1 *. . . I*, we use the model:

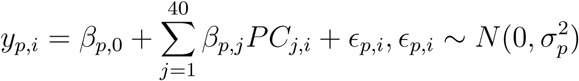

We visualize the data in Figure 5 with the values [inline1] For UKB figures we varied the number of input PCs (5 *. . .* 40) into UMAP as well as the UMAP parameters (setting the number of neighbours to 5 or 10, the minimum distance to 0 or 0.01, and the dimensionality to 3 or 5) and set the HDBSCAN(*E*^) parameters to 25 minimum points and *E*^to 0.5. This resulted in 36 × 2 × 2 × 2 = 288 unique runs of parameters. To test the robustness of the clustering (shown in Figure S14), we re-ran a subset of the parametrizations for a total of 604 runs.

## SI Figures

**Figure S1:**
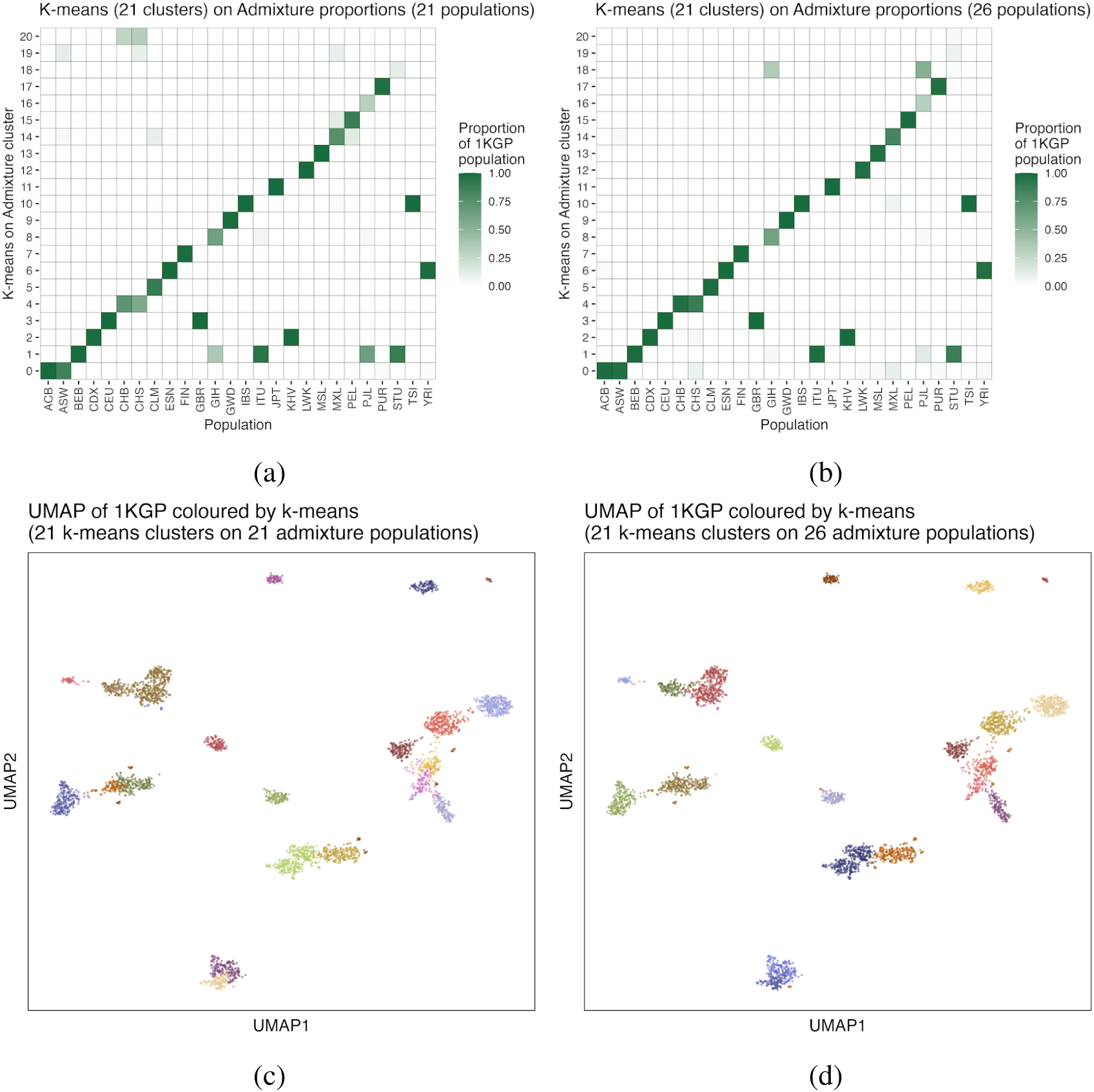
Clustering 1KGP data using ADMIXTURE 1.3.0. (a) Generating 21 clusters using k-means clustering on admixture proportions (*K* = 21 populations specified). The adjusted Rand Index compared to population labels is 0.611. (b) Generating 21 clusters using k-means clustering on admixture proportions (*K* = 26 populations specified). The adjusted Rand Index compared to population labels is 0.661. Using density clustering gives an adjusted Rand Index of 0.769. (c) UMAP of the 1KGP coloured by k-means clustering of admixture proportions (21 clusters on *K* = 21 admixture populations). (c) UMAP of the 1KGP coloured by k-means clustering of admixture proportions (21 clusters on *K* = 26 admixture populations).

**Figure S2:**
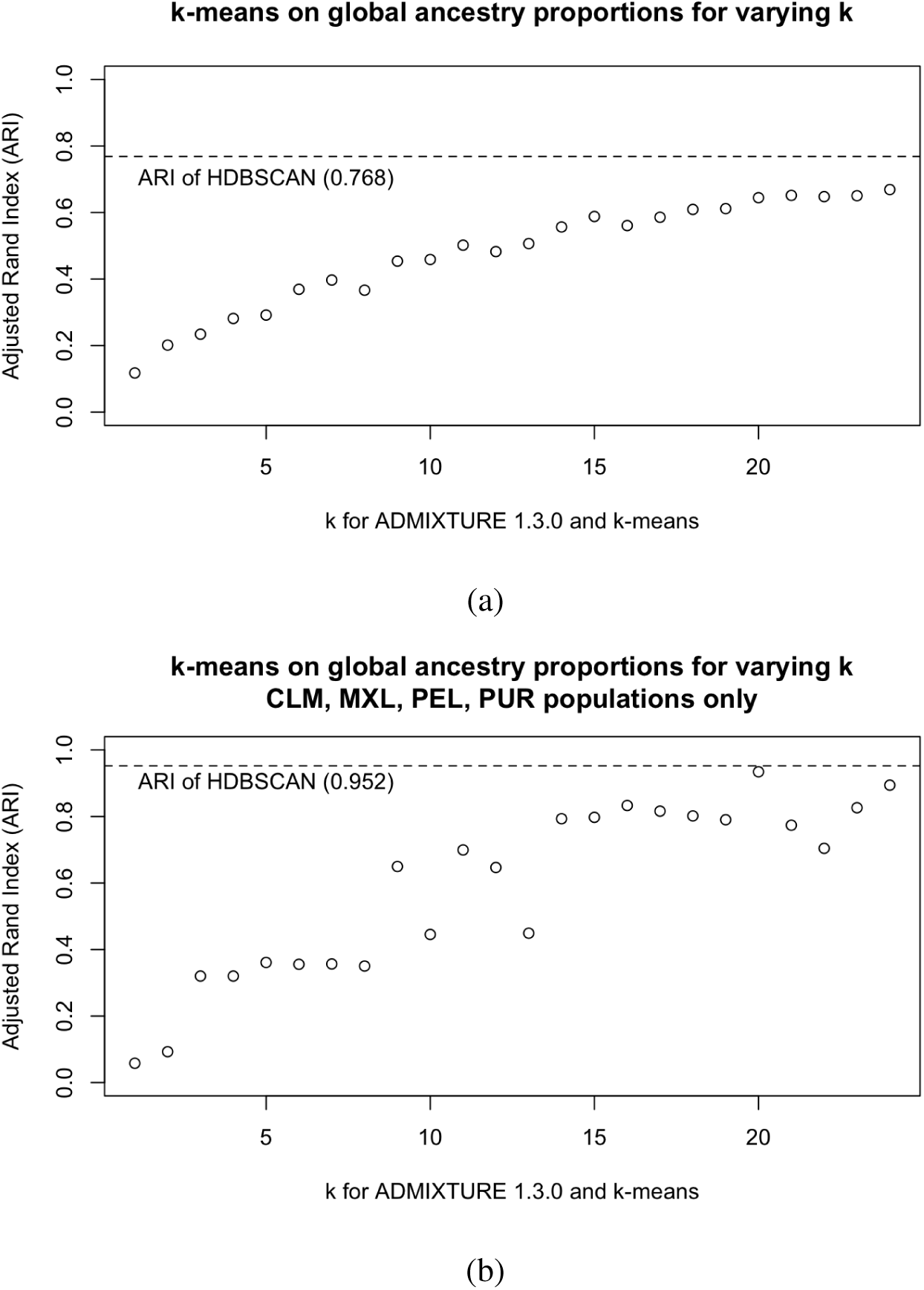
Adjusted Rand Indices (ARI) comparing hard clustering of global ancestry estimates. We generated ancestry proportions from 1KGP individuals assuming *k* = {3, *. . .*, 26} source populations, and then ran k-means on the individual-level proportions, assuming a corresponding value of *k* clusters (e.g. for ADMIXTURE with *k* = 10 we also ran k-means assuming 10 clusters). We use 1KGP population labels as ground truth, and the ARI of HDBSCAN illustrated in Figure 2b is presented for comparison. An ARI closer to 1 is considered closer to ground truth. Top: ARI for all 1KGP populations. Bottom: ARI for the CLM, MXL, PEL, PUR populations.

**Figure S3:**
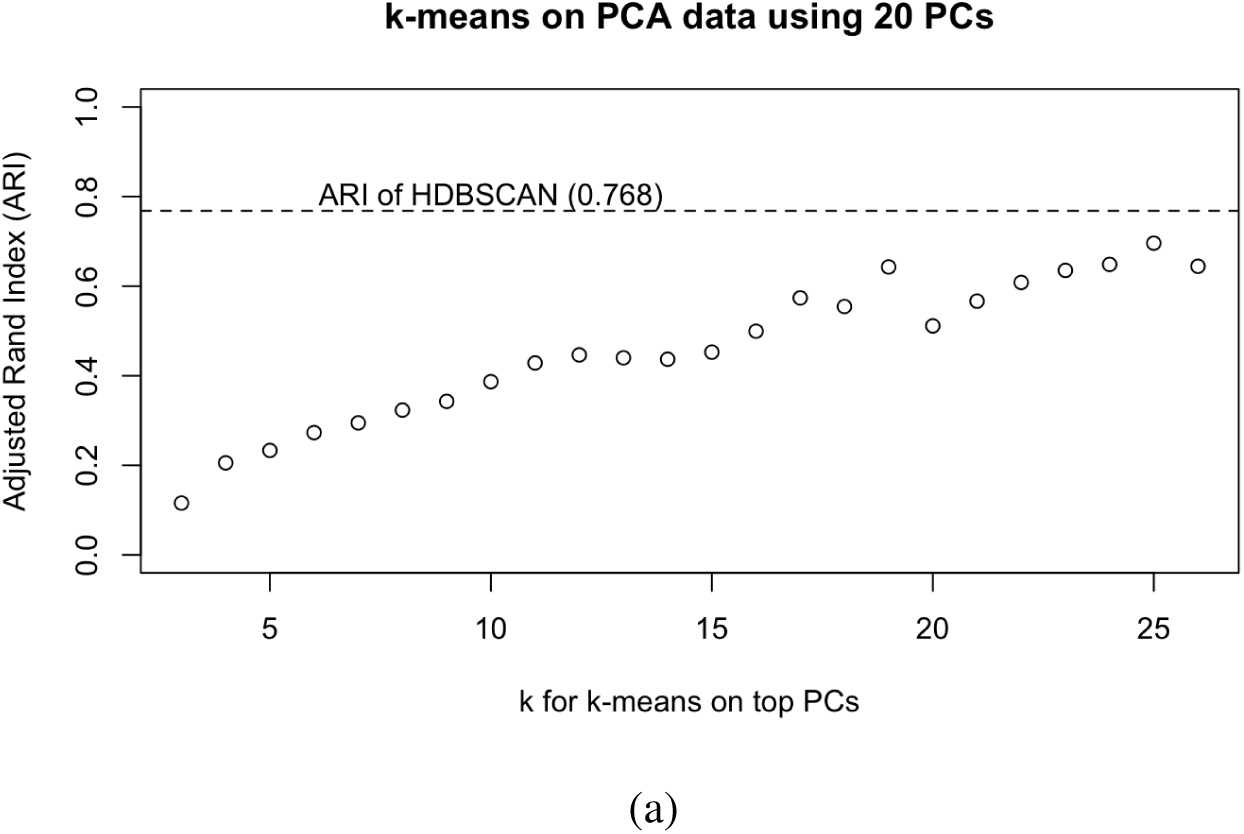
Adjusted Rand Indices (ARI) comparing k-means clustering of the top 20 PCs of the 1KGP data. We applied k-means clustering assuming *k* = 3 26. We use 1KGP population labels as ground truth, and the ARI of HDBSCAN illustrated in Figure 2b is presented for comparison. An ARI closer to 1 is considered closer to ground truth. We also applied k-means clustering to the top 5, 10, 15, 25, 30, 35, 40, 45, 50 PCs but the results were similar. We illustrate 20 PCs specifically because it provided the highest ARI (0.696 at *k* = 25).

**Table S1:**
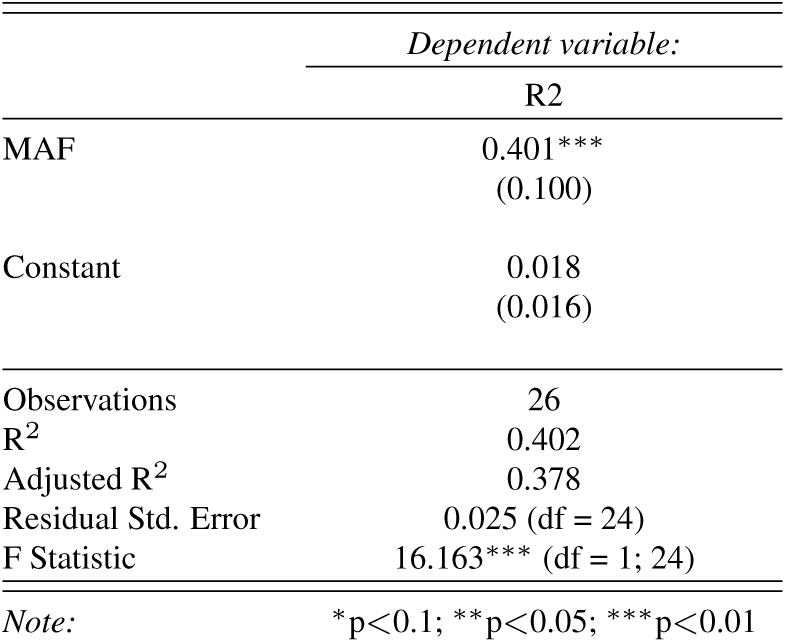
Linear regression model between minor allele frequency (MAF) of *rs*4420638 within each cluster from Figure 4a and the *R*^2^ of a PGS for LDL generated by VIPRS using the clusters from Figure 4a. The plot of the regression is present in Figure S12.

**Figure S4:**
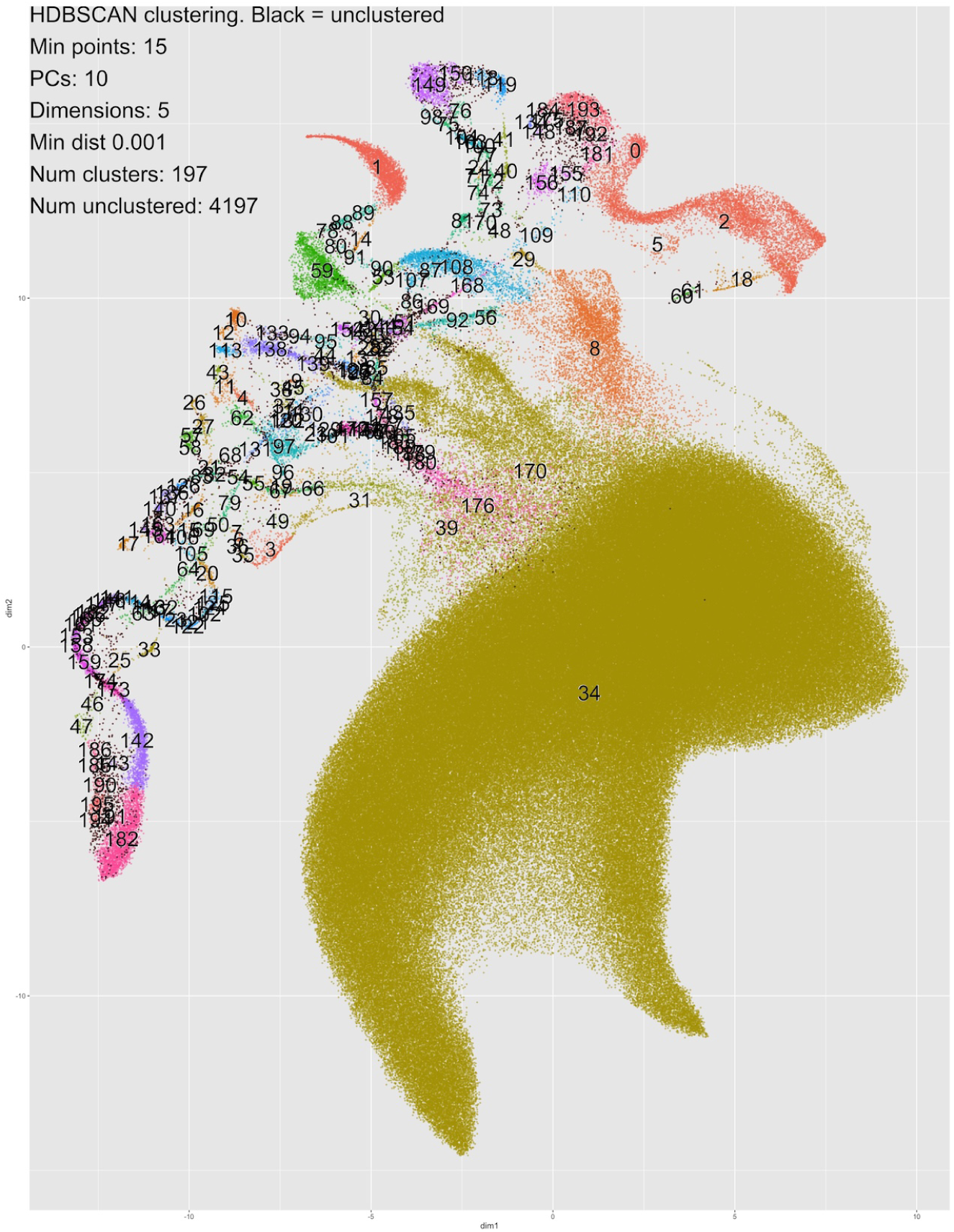
An example of a clustering of the UKB data using HDBSCAN rather than HDBSCAN(*E*^). The algorithm fails to adequately cluster many of the sub-populations, categorizing 4, 197 individuals as noise and generated almost 200 micro-clusters.

**Figure S5:**
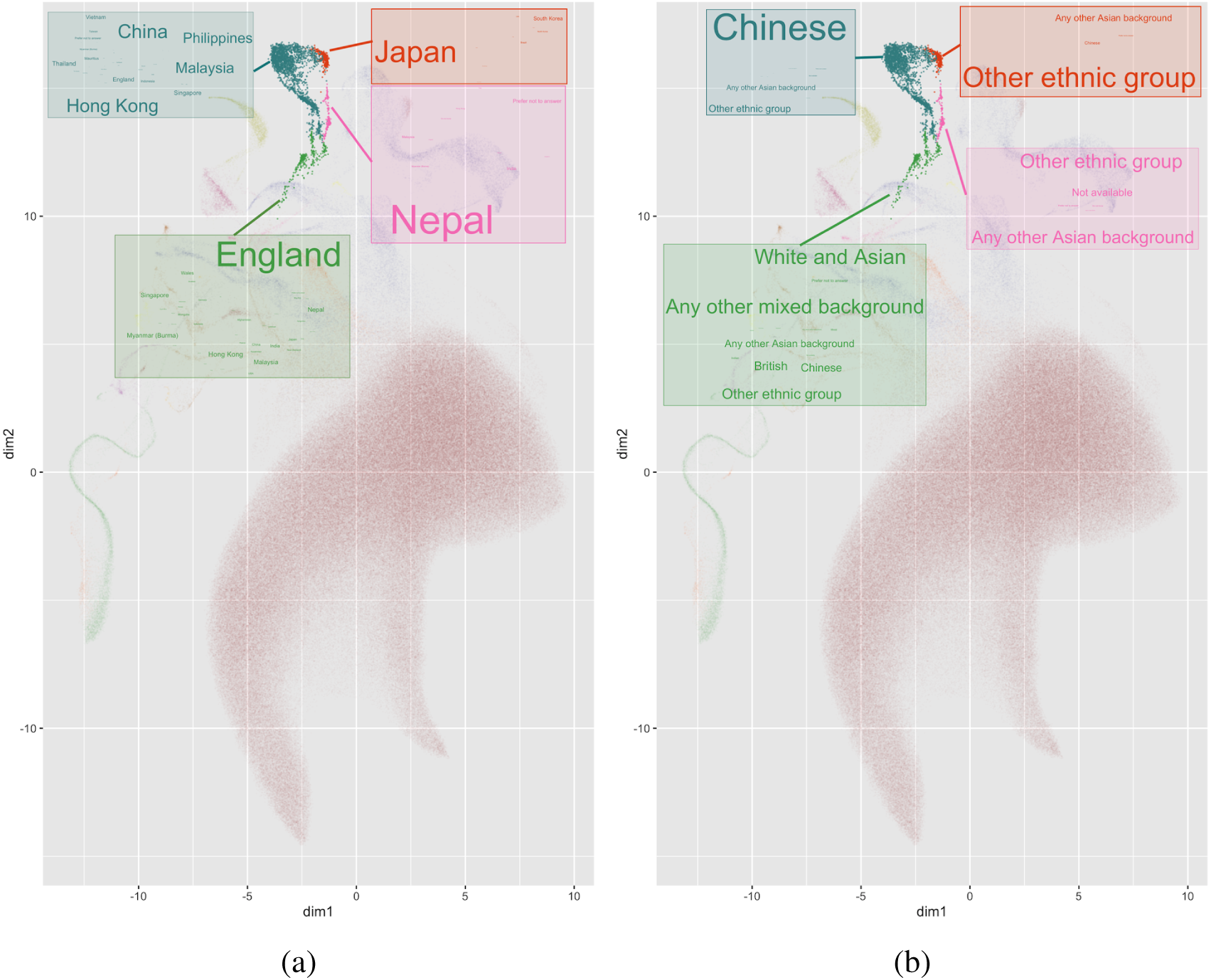
Word clouds generated from four clusters in the UKB from Figure 4. (a) Left: Word clouds of the most common countries of birth within each cluster. Most individuals in the orange cluster (Cluster 0) were born in Japan, and most in the pink cluster (Cluster 15) were born in Nepal. (b) Right: Word clouds for the most common EB. The most common in the blue cluster (Cluster 13) was “Chinese”, while those in the green cluster (Cluster 14) select a variety, including “White British”, “Chinese”, “Mixed”, or “Other”. Detailed breakdowns are available in Tables S9 and S10.

**Figure S6:**
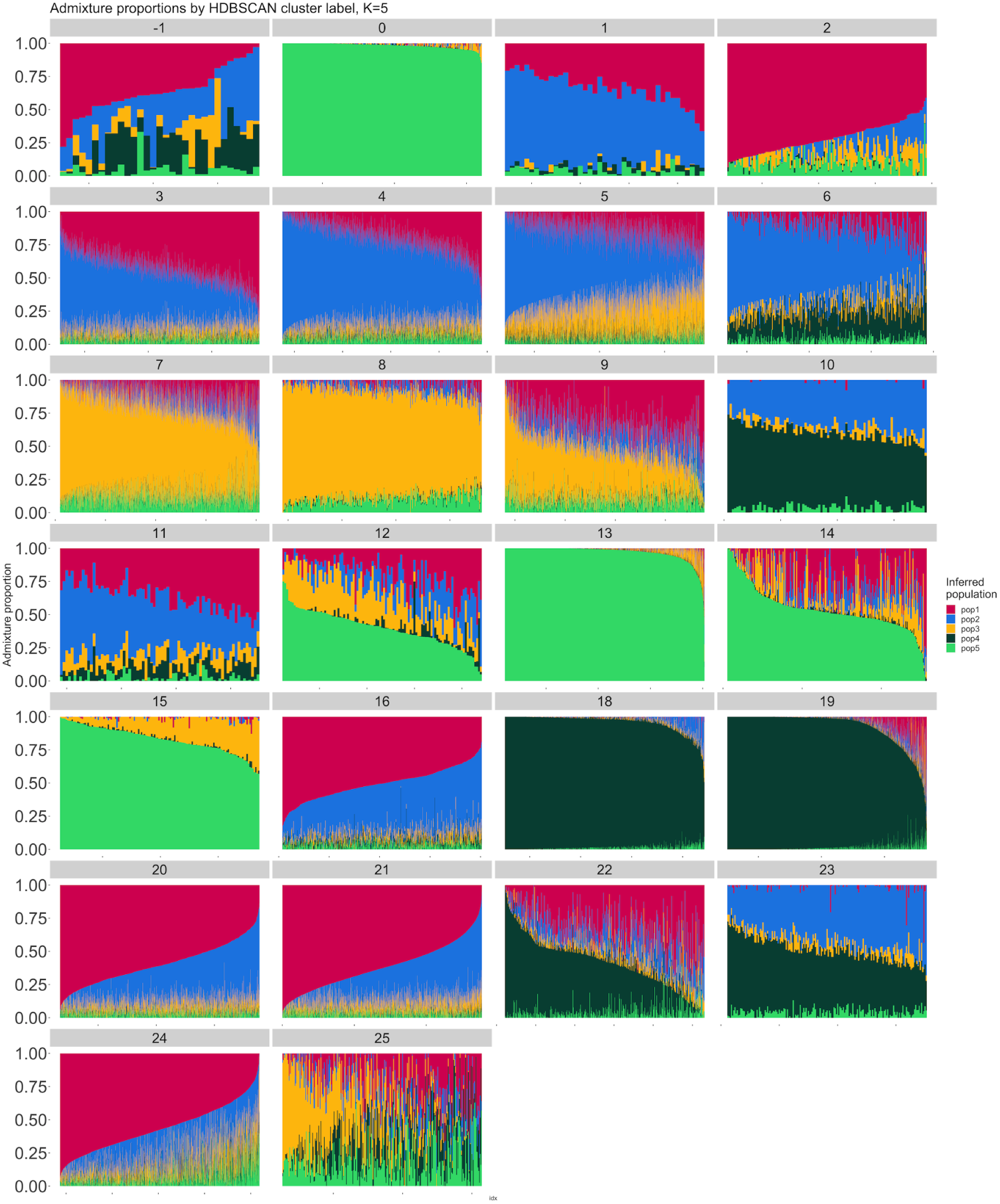
Admixture proportions for *K* = 5 populations on each of the clusters in Figure 4. Cluster 17 (*n >* 400, 000) was excluded for computational reasons. Individuals not assigned to a cluster are labelled as −1.

**Figure S7:**
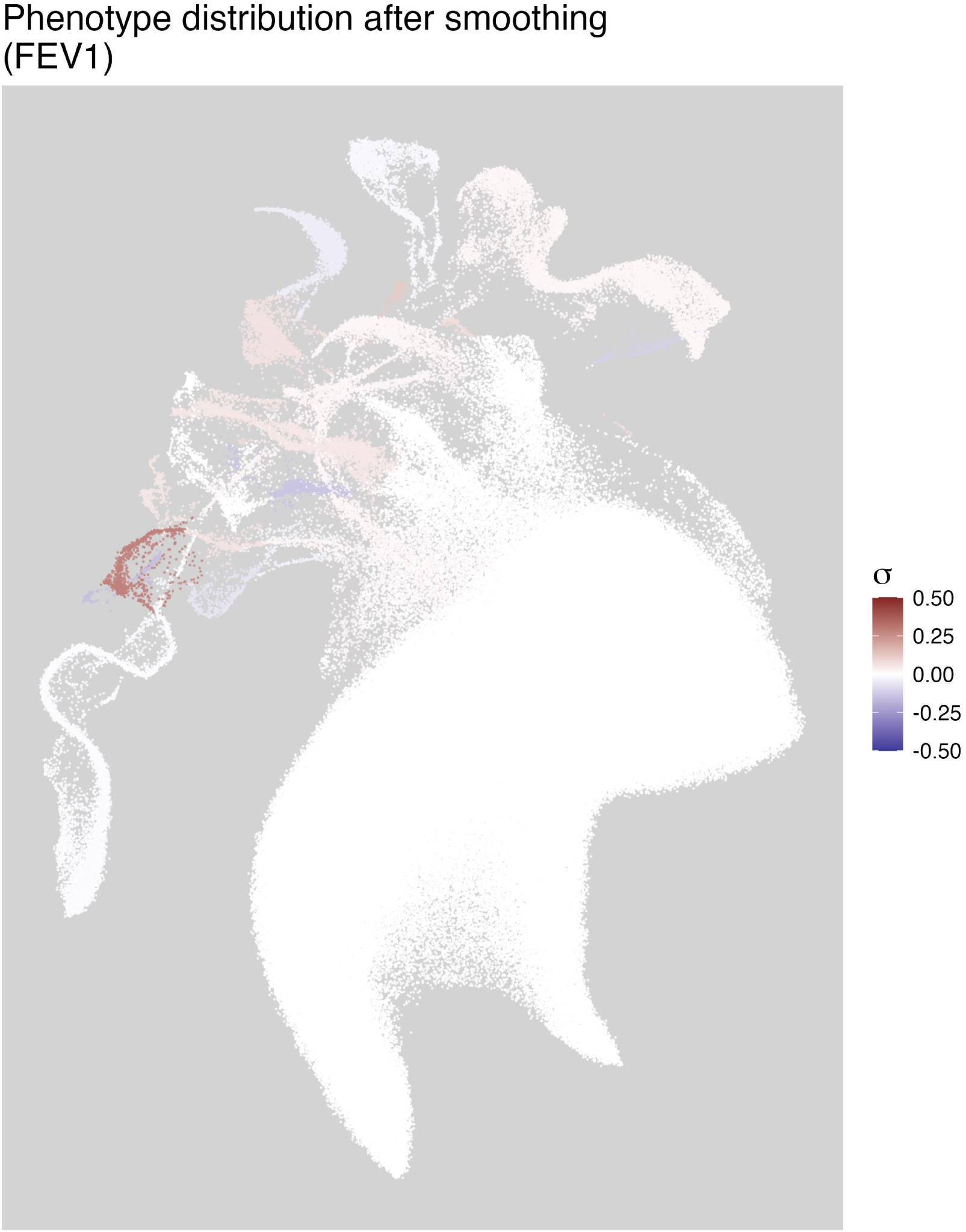
FEV1 values averaged by a single run of clustering rather than smoothed over multiple runs of clustering. Patterns that appear in Figure 5 are obscured because, e.g., smaller clusters may have been merged into larger ones in this particular set of parameters.

**Figure S8:**
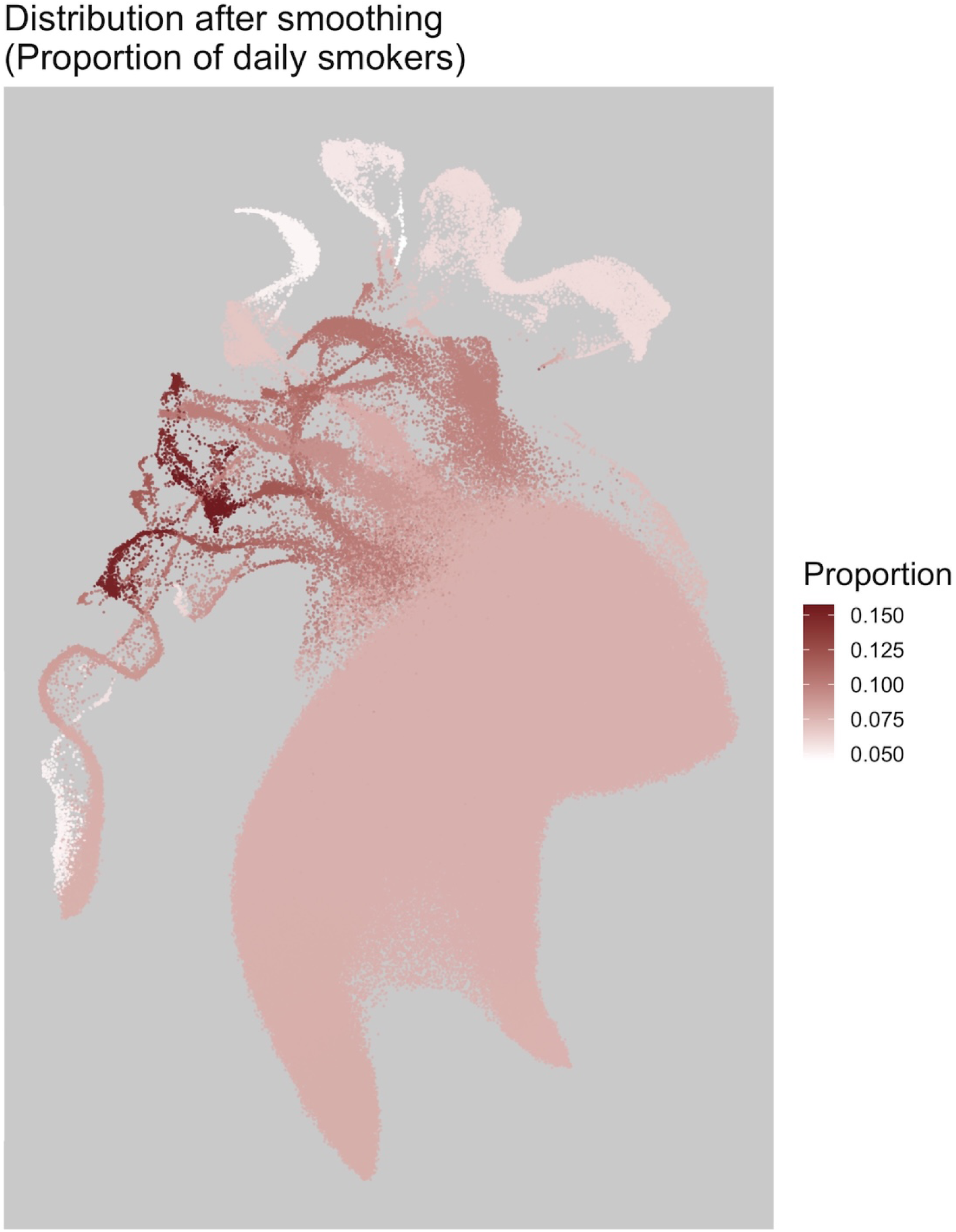
Proportion of daily smokers, smoothed using Algorithm 1.

**Figure S9:**
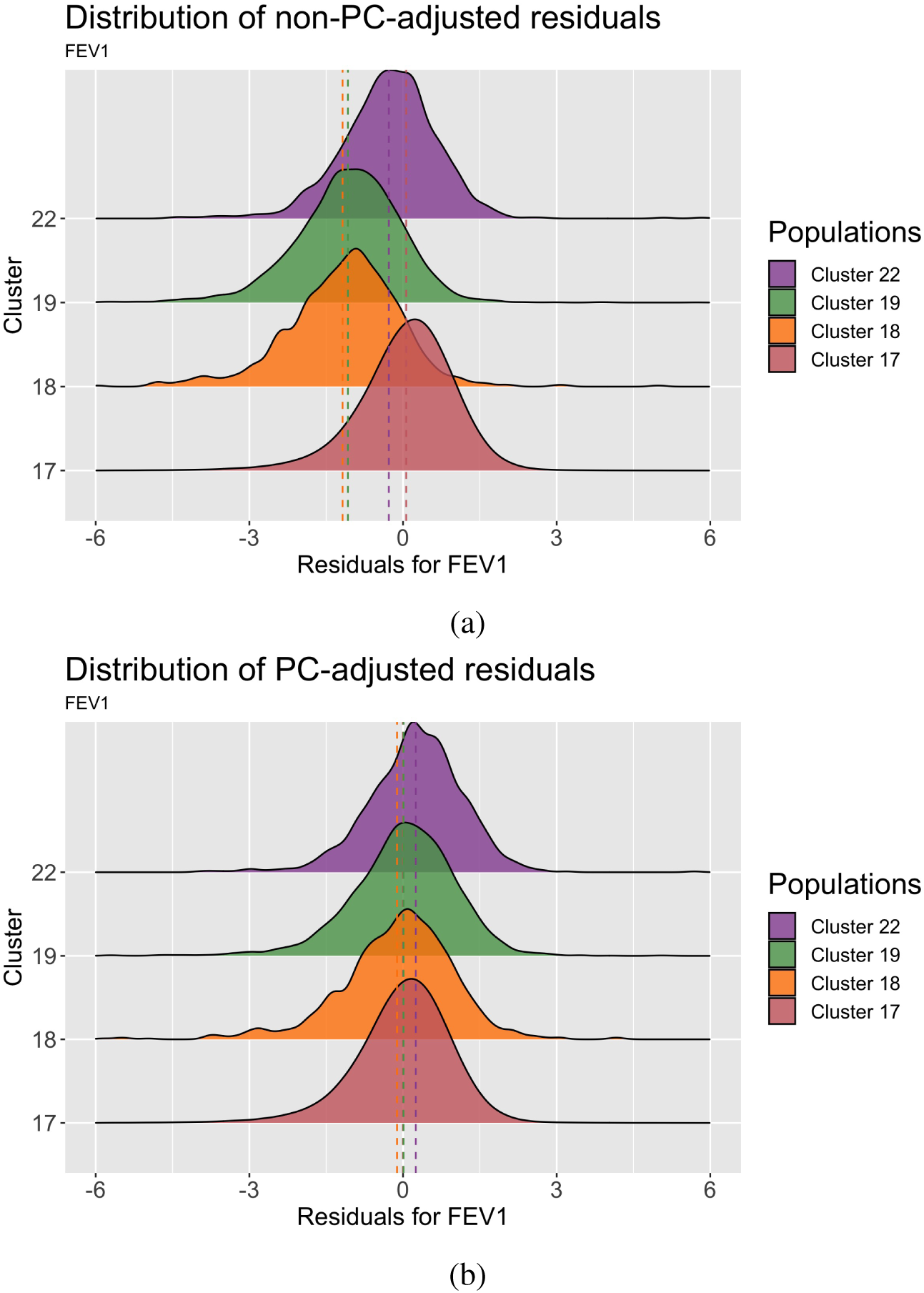
Distributions of FEV1 adjusted for age and sex stratified by cluster. Vertical dotted lines represent the mean of the distribution. Cluster labels and colours match those in Figure 4a. Cluster 17 is mostly European-born individuals, Cluster 18 is mostly sub-Saharan African born individuals, Cluster 19 is mostly individuals born in England, the Caribbean, Ghana, and Nigeria, and Cluster 22 is mostly individuals born in England who chose the EB “White and Black Caribbean” or “White and Black African”. (a) Top: Distribution of FEV1 by cluster without adjusting for population structure. (b) Bottom: Distribution of FEV1 by cluster after having adjusted for the top 40 principal components.

**Figure S10:**
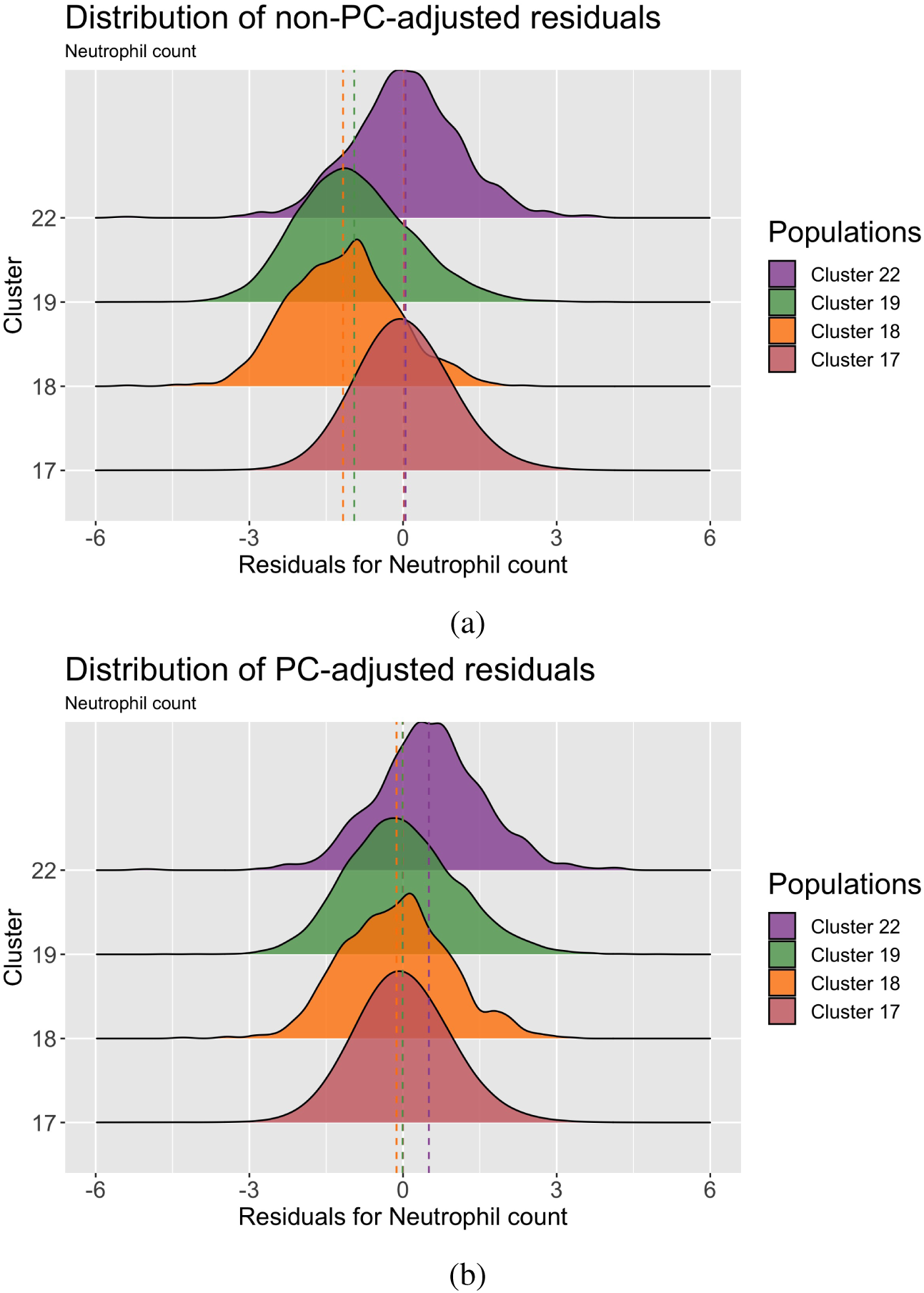
Distributions of neutrophil count adjusted for age and sex stratified by cluster. Vertical dotted lines represent the mean of the distribution. Cluster labels and colours match those in Figure 4a. Cluster 17 is mostly European-born individuals, Cluster 18 is mostly sub-Saharan African born individuals, Cluster 19 is mostly individuals born in England, the Caribbean, Ghana, and Nigeria, and Cluster 22 is mostly individuals born in England who chose the EB “White and Black Caribbean” or “White and Black African”. (a) Top: Distribution of neutrophil count by cluster without adjusting for population structure. (b) Bottom: Distribution of neutrophil count by cluster after having adjusted for the top 40 principal components.

**Figure S11:**
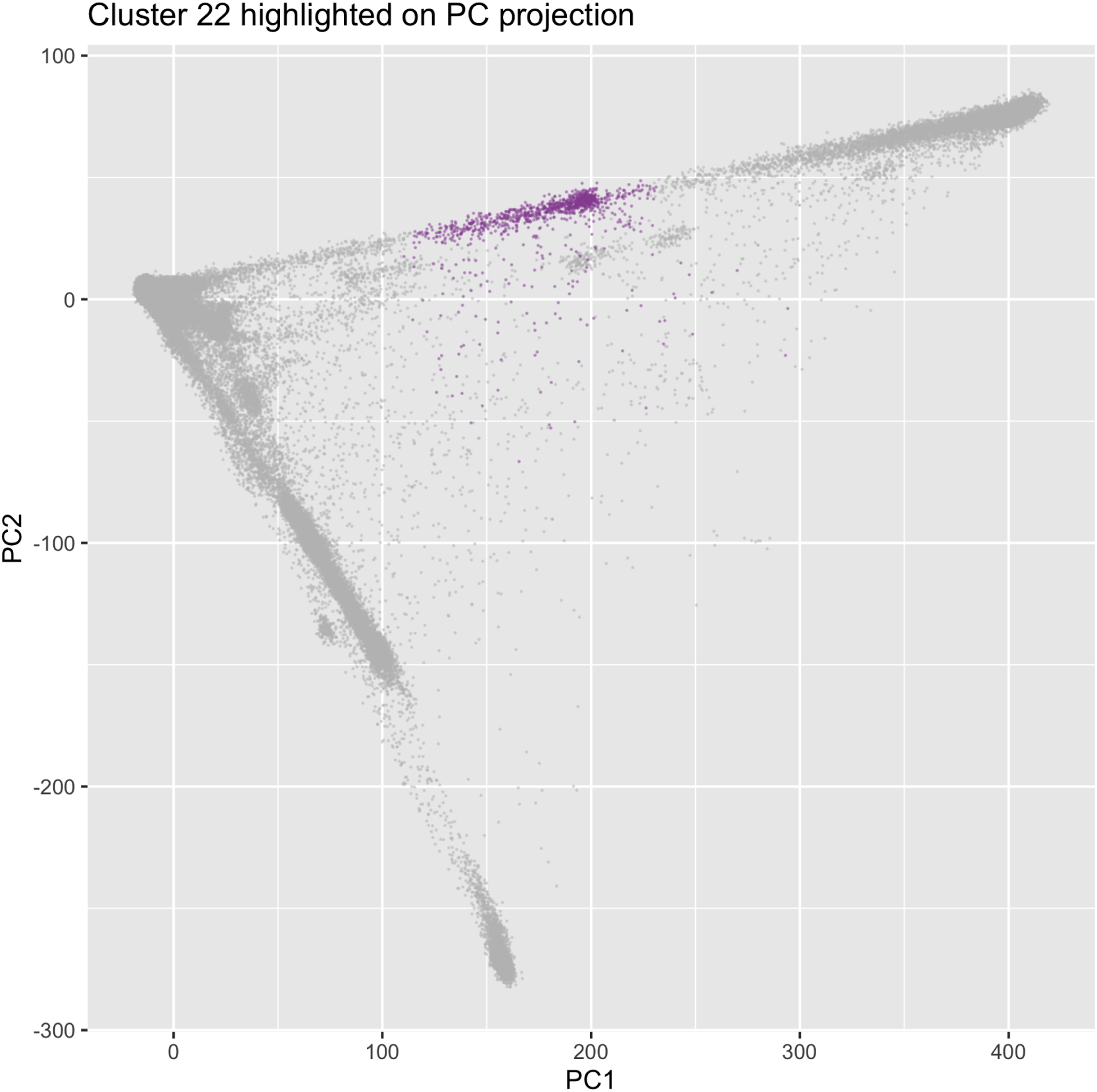
Cluster 22 from Figure 4a highlighted coloured in on a plot of PC1 and PC2.

**Figure S12:**
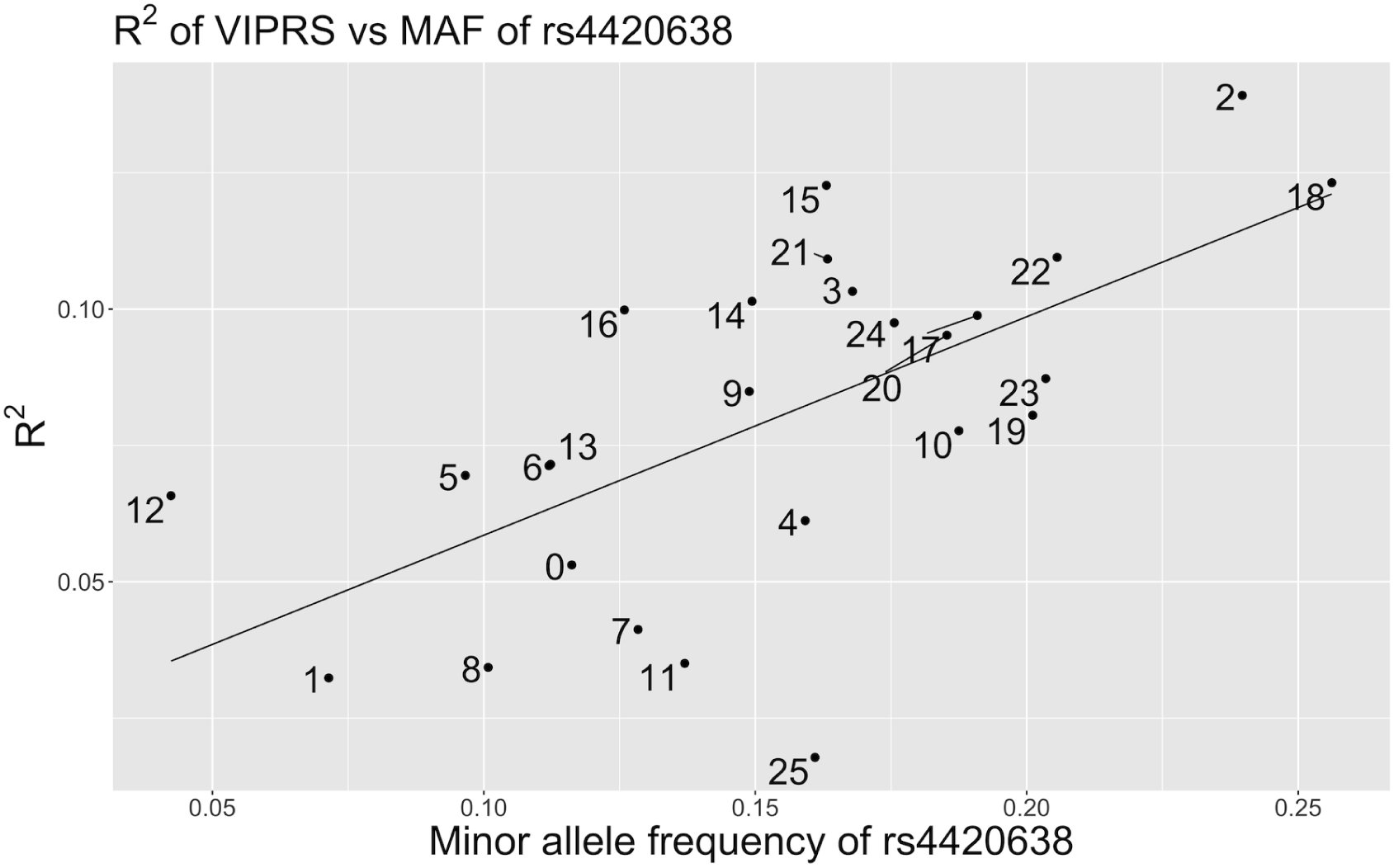
Regression line of the *R*^2^ of a PGS generated by VIPRS versus the minor allele frequency *rs*4420638, labelled by clusters from Figure 4a. The regression summary is presented in Table S1.

**Table S2:** Linear regression model between minor allele frequency (MAF) of *rs*7412 within each cluster from Figure 4a and the *R*^2^ of a PGS for LDL generated by VIPRS using the clusters from Figure 4a. The plot of the regression is present in Figure S13.

| *Dependent variable:* |  |
| --- | --- |
| R2 |  |
| MAF | 0.609***<br>(0.202) |
| Constant | 0.041***<br>(0.014) |
| Observations | 26 |
| R<sup>2</sup> | 0.275 |
| Adjusted R<sup>2</sup> | 0.245 |
| Residual Std. Error | 0.027 (df = 24) |
| F Statistic | 9.126*** (df = 1; 24) |
| *Note:* *p<0.1; **p<0.05; ***p<0.01 |  |

**Figure S13:**
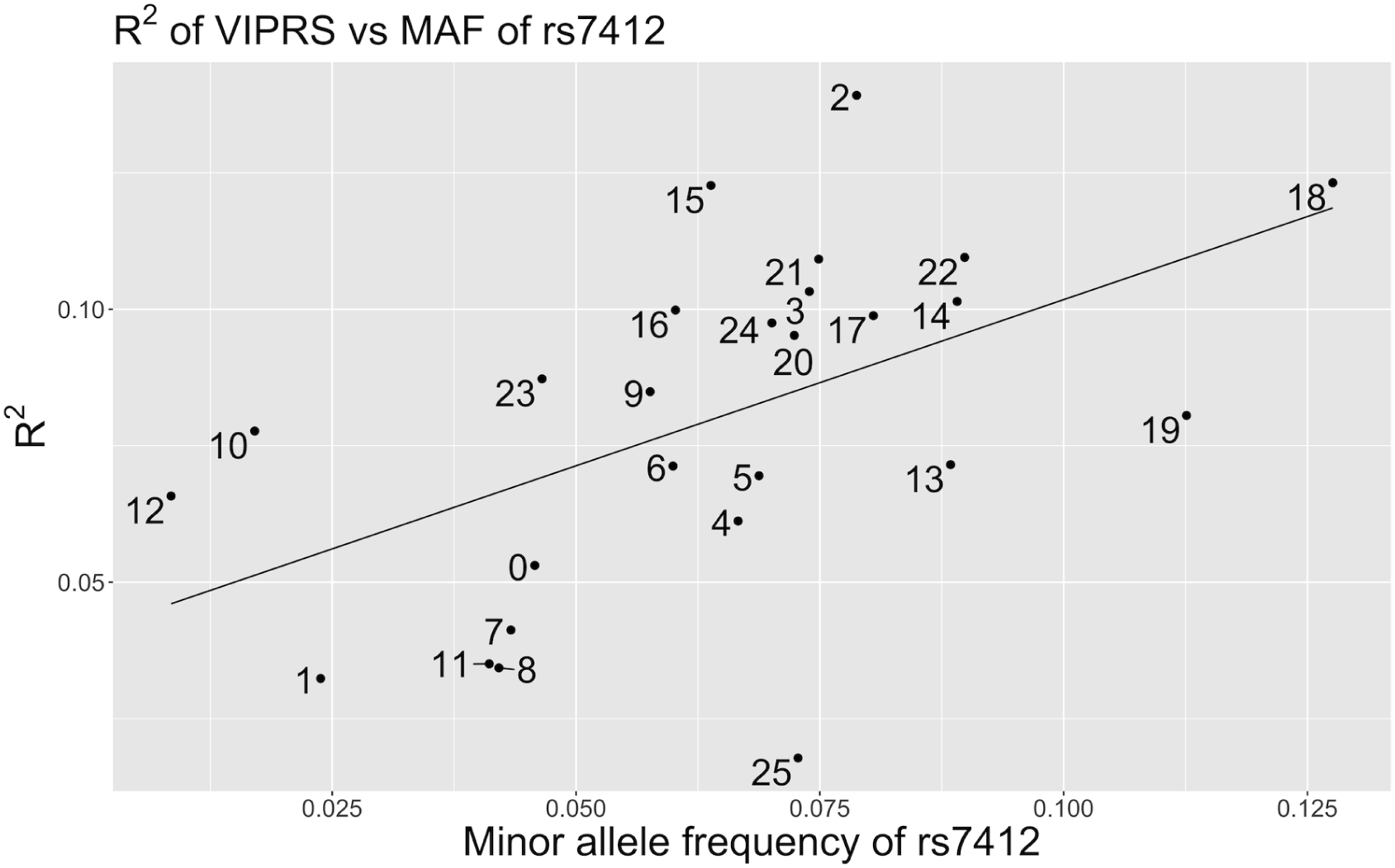
Regression line of the *R*^2^ of a PGS generated by VIPRS versus the minor allele frequency *rs*7412, labelled by clusters from Figure 4a. The regression summary is presented in Table S2.

**Table S3:** HDBSCAN run on PCA data alone classifies many individuals as noise. Minimum points were set to 5, *E*^ = 05

| Number PCs | Unclustered individuals |
| --- | --- |
| 2 | 8970 |
| 3 | 19701 |
| 4 | 61938 |
| 5 | 11423 |
| 6 | 92124 |
| 7 | 99824 |
| 8 | 65442 |
| 9 | 113299 |

**Figure S14:**
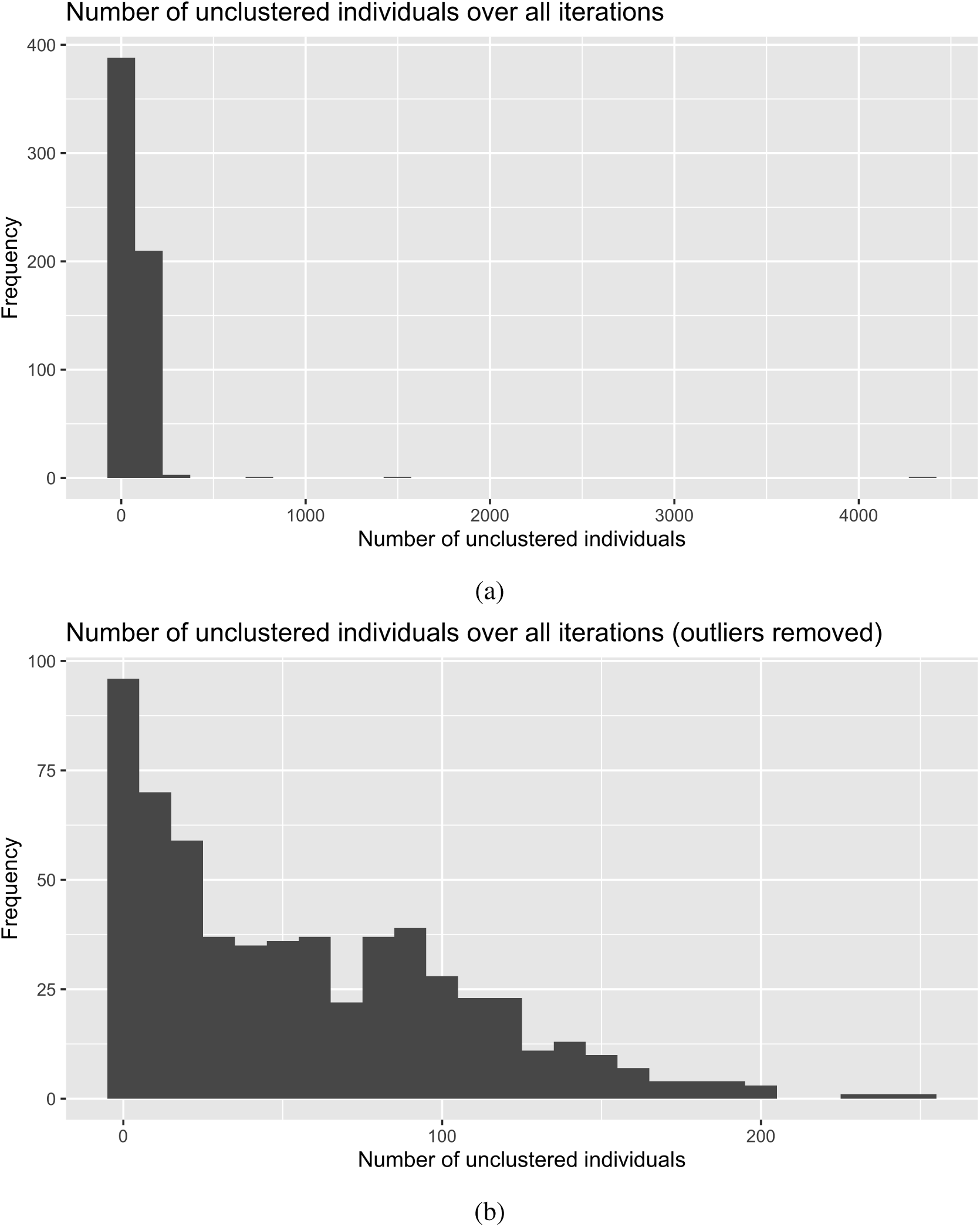
For 604 runs of UMAP-HDBSCAN(*E*^) on the UKB, we count the number of individuals not assigned to a cluster. (a) Top: Across all 604 runs. (b) Bottom: To improve the scale of the figure, we remove 3 outlier runs in which 684, 1, 535, and 4, 346 individuals were not assigned to a cluster.

**Figure S15:**
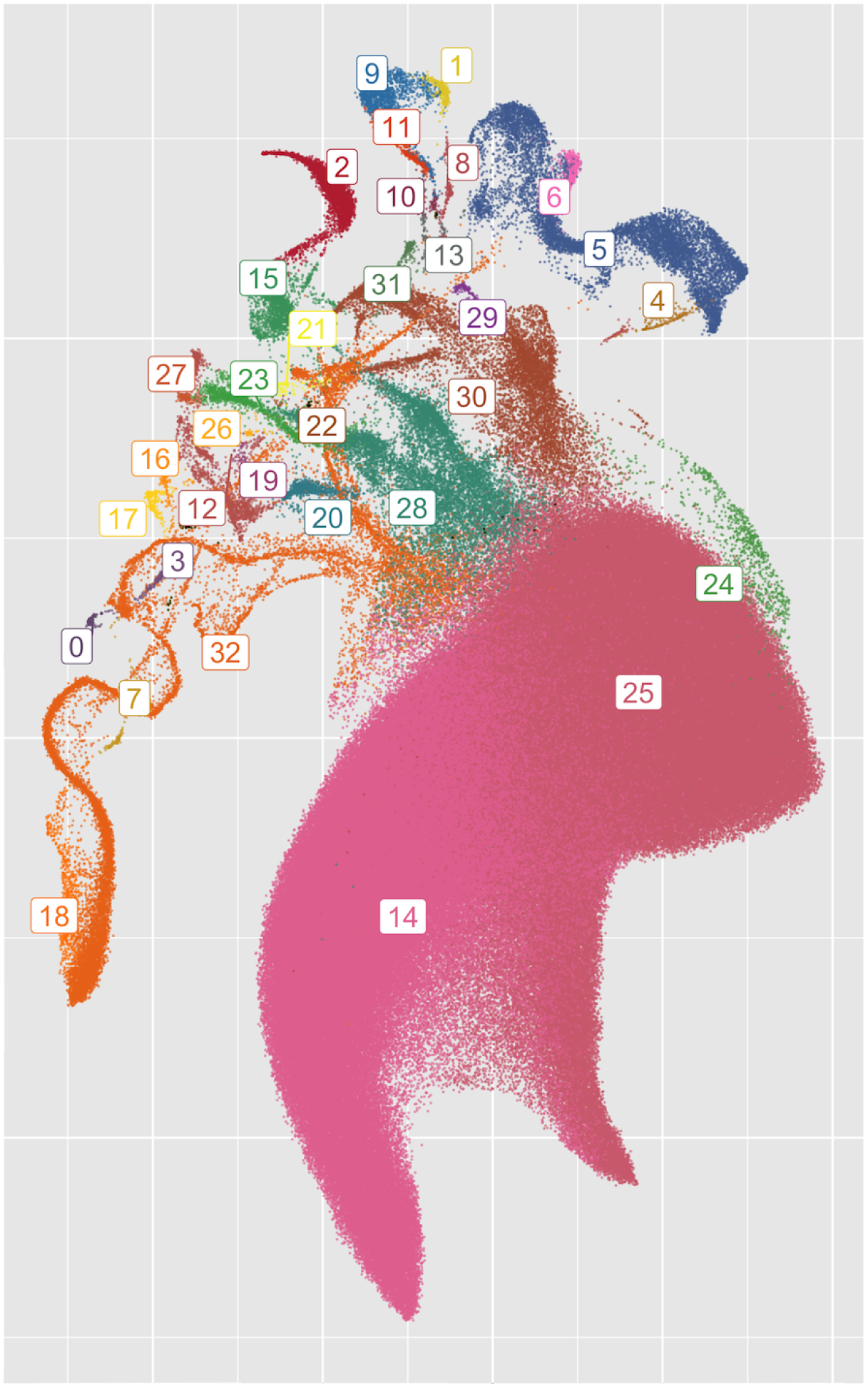
An alternative clustering of UKB data. Compared to Figure 4a, the largest cluster (Cluster 17 in that figure) has been split into three smaller clusters (Clusters 14, 24, 25 in this figure). Other clusters have been split or merged, while some remain the same between runs.

**Table S4:** Names and abbreviations of 1KGP populations.

| Abbreviation | Population name |
| --- | --- |
| ACB | African Caribbean in Barbados |
| ASW | African Ancestry in SW USA |
| BEB | Bengali in Bangladesh |
| CDX | Chinese Dai in Xishuangbanna, China |
| CEU | Utah residents with Northern/Western European ancestry |
| CHB | Han Chinese in Beijing, China |
| CHS | Han Chinese South |
| CLM | Colombian in Medellín, Colombia |
| ESN | Esan in Nigeria |
| FIN | Finnish in Finland |
| GBR | British From England and Scotland |
| GWD | Gambian in Western Division – Mandinka |
| GIH | Gujarati Indians in Houston, Texas, USA |
| IBS | Iberian Populations in Spain |
| ITU | Indian Telugu in the UK |
| JPT | Japanese in Tokyo, Japan |
| KHV | Kinh in Ho Chi Minh City, Vietnam |
| LWK | Luhya in Webuye, Kenya |
| MSL | Mende in Sierra Leone |
| MXL | Mexican Ancestry in Los Angeles, CA, USA |
| PEL | Peruvian in Lima, Peru |
| PJL | Punjabi in Lahore, Pakistan |
| PUR | Puerto Rican in Puerto Rico |
| STU | Sri Lankan Tamil in the UK |
| TSI | Toscani in Italy |
| YRI | Yoruba in Ibadan, Nigeria |

**Table S5:** Cluster assignments for each 1KGP population, showing how many individuals from each population ended up in each cluster.

| 1KGP population | Cluster label | 1KGP in cluster | Total in 1KGP | Proportion in cluster |
| --- | --- | --- | --- | --- |
| ACB | 0 | 122 | 122 | 1.0000000 |
| ASW | 0 | 103 | 107 | 0.9626168 |
| ASW | 5 | 3 | 107 | 0.0280374 |
| ASW | 15 | 1 | 107 | 0.0093458 |
| BEB | 1 | 133 | 143 | 0.9300699 |
| BEB | 11 | 10 | 143 | 0.0699301 |
| CDX | 2 | 104 | 105 | 0.9904762 |
| CDX | 4 | 1 | 105 | 0.0095238 |
| CEU | 3 | 183 | 183 | 1.0000000 |
| CHB | 4 | 105 | 105 | 1.0000000 |
| CHS | 4 | 171 | 171 | 1.0000000 |
| CLM | 5 | 142 | 146 | 0.9726027 |
| CLM | 15 | 4 | 146 | 0.0273973 |
| ESN | 6 | 172 | 172 | 1.0000000 |
| FIN | 7 | 104 | 104 | 1.0000000 |
| GBR | 3 | 105 | 105 | 1.0000000 |
| GIH | 8 | 69 | 111 | 0.6216216 |
| GIH | 17 | 41 | 111 | 0.3693694 |
| GIH | 11 | 1 | 111 | 0.0090090 |
| GWD | 9 | 179 | 180 | 0.9944444 |
| GWD | 14 | 1 | 180 | 0.0055556 |
| IBS | 10 | 162 | 162 | 1.0000000 |
| ITU | 11 | 109 | 118 | 0.9237288 |
| ITU | 17 | 9 | 118 | 0.0762712 |
| JPT | 12 | 104 | 105 | 0.9904762 |
| JPT | 4 | 1 | 105 | 0.0095238 |
| KHV | 2 | 118 | 121 | 0.9752066 |
| KHV | 4 | 3 | 121 | 0.0247934 |
| LWK | 13 | 110 | 110 | 1.0000000 |
| MSL | 14 | 122 | 122 | 1.0000000 |
| MXL | 15 | 97 | 104 | 0.9326923 |
| MXL | 10 | 7 | 104 | 0.0673077 |
| PEL | 16 | 128 | 129 | 0.9922481 |
| PEL | 5 | 1 | 129 | 0.0077519 |
| PJL | 17 | 95 | 155 | 0.6129032 |
| PJL | 19 | 48 | 155 | 0.3096774 |
| PJL | 11 | 12 | 155 | 0.0774194 |
| PUR | 18 | 145 | 149 | 0.9731544 |
| PUR | 0 | 4 | 149 | 0.0268456 |
| STU | 11 | 124 | 128 | 0.9687500 |
| STU | 17 | 4 | 128 | 0.0312500 |
| TSI | 10 | 111 | 111 | 1.0000000 |
| YRI | 20 | 181 | 182 | 0.9945055 |
| YRI | 6 | 1 | 182 | 0.0054945 |

**Table S6:** Composition of each cluster broken down by 1KGP population.

| Cluster | 1KGP population | 1KGP population in cluster | Proportion |
| --- | --- | --- | --- |
| 0 | ACB | 122 | 0.5327511 |
| 0 | ASW | 103 | 0.4497817 |
| 0 | PUR | 4 | 0.0174672 |
| 1 | BEB | 133 | 1.0000000 |
| 2 | CDX | 104 | 0.4684685 |
| 2 | KHV | 118 | 0.5315315 |
| 3 | CEU | 183 | 0.6354167 |
| 3 | GBR | 105 | 0.3645833 |
| 4 | CDX | 1 | 0.0035587 |
| 4 | CHB | 105 | 0.3736655 |
| 4 | CHS | 171 | 0.6085409 |
| 4 | JPT | 1 | 0.0035587 |
| 4 | KHV | 3 | 0.0106762 |
| 5 | ASW | 3 | 0.0205479 |
| 5 | CLM | 142 | 0.9726027 |
| 5 | PEL | 1 | 0.0068493 |
| 6 | ESN | 172 | 0.9942197 |
| 6 | YRI | 1 | 0.0057803 |
| 7 | FIN | 104 | 1.0000000 |
| 8 | GIH | 69 | 1.0000000 |
| 9 | GWD | 179 | 1.0000000 |
| 10 | IBS | 162 | 0.5785714 |
| 10 | MXL | 7 | 0.0250000 |
| 10 | TSI | 111 | 0.3964286 |
| 11 | BEB | 10 | 0.0390625 |
| 11 | GIH | 1 | 0.0039062 |
| 11 | ITU | 109 | 0.4257812 |
| 11 | PJL | 12 | 0.0468750 |
| 11 | STU | 124 | 0.4843750 |
| 12 | JPT | 104 | 1.0000000 |
| 13 | LWK | 110 | 1.0000000 |
| 14 | GWD | 1 | 0.0081301 |
| 14 | MSL | 122 | 0.9918699 |
| 15 | ASW | 1 | 0.0098039 |
| 15 | CLM | 4 | 0.0392157 |
| 15 | MXL | 97 | 0.9509804 |
| 16 | PEL | 128 | 1.0000000 |
| 17 | GIH | 41 | 0.2751678 |
| 17 | ITU | 9 | 0.0604027 |
| 17 | PJL | 95 | 0.6375839 |
| 17 | STU | 4 | 0.0268456 |
| 18 | PUR | 145 | 1.0000000 |
| 19 | PJL | 48 | 1.0000000 |
| 20 | YRI | 181 | 1.0000000 |

**Table S7:** HDBSCAN(*E*^) carried out on the top PCs of the UKB for varying values of the minimum number of points, *E*^, and the number of input PCs. Regardless of parameters, HDBSCAN is unable to cluster many individuals in the UKB without UMAP as an intermediate step.

| Min. points | $\hat{\epsilon}$ | Number PCs | Number unclustered | Min. points | $\hat{\epsilon}$ | Number PCs | Number unclustered |
| --- | --- | --- | --- | --- | --- | --- | --- |
| 5 | 0 | 2 | 218613 | 10 | 0 | 2 | 273869 |
| 5 | 0 | 3 | 317131 | 10 | 0 | 3 | 14369 |
| 5 | 0 | 4 | 384120 | 10 | 0 | 4 | 16778 |
| 5 | 0 | 5 | 11423 | 10 | 0 | 5 | 63767 |
| 5 | 0 | 6 | 92124 | 10 | 0 | 6 | 52090 |
| 5 | 0 | 7 | 99824 | 10 | 0 | 7 | 52906 |
| 5 | 0 | 8 | 65442 | 10 | 0 | 8 | 45236 |
| 5 | 0 | 9 | 113299 | 10 | 0 | 9 | 66286 |
| 5 | 0.3 | 2 | 14217 | 10 | 0.3 | 2 | 11900 |
| 5 | 0.3 | 3 | 30645 | 10 | 0.3 | 3 | 14369 |
| 5 | 0.3 | 4 | 183491 | 10 | 0.3 | 4 | 16778 |
| 5 | 0.3 | 5 | 11423 | 10 | 0.3 | 5 | 63767 |
| 5 | 0.3 | 6 | 92124 | 10 | 0.3 | 6 | 52090 |
| 5 | 0.3 | 7 | 99824 | 10 | 0.3 | 7 | 52906 |
| 5 | 0.3 | 8 | 65442 | 10 | 0.3 | 8 | 45236 |
| 5 | 0.3 | 9 | 113299 | 10 | 0.3 | 9 | 66286 |
| 5 | 0.5 | 2 | 8970 | 10 | 0.5 | 2 | 8972 |
| 5 | 0.5 | 3 | 19701 | 10 | 0.5 | 3 | 14369 |
| 5 | 0.5 | 4 | 61938 | 10 | 0.5 | 4 | 16778 |
| 5 | 0.5 | 5 | 11423 | 10 | 0.5 | 5 | 63767 |
| 5 | 0.5 | 6 | 92124 | 10 | 0.5 | 6 | 52090 |
| 5 | 0.5 | 7 | 99824 | 10 | 0.5 | 7 | 52906 |
| 5 | 0.5 | 8 | 65442 | 10 | 0.5 | 8 | 45236 |
| 5 | 0.5 | 9 | 113299 | 10 | 0.5 | 9 | 66286 |

**Table S8:** Possible values for ethnic background in the UKB (Data-Field 21000). Participants are first asked “What is your ethnic group?” and then asked “What is your ethnic background?” For “Chinese”, there is no second question. Participants may also select “Prefer not to answer” for the second question; it is possible to have ethnic background recorded as ethnic group (e.g. just “White” or “Mixed”. Excluding “Do not know”, “Prefer not to answer”, and “Not available”, there were 20 unique values of ethnic background.

| Ethnic group | Ethnic background |
| --- | --- |
| White | British |
| White | Irish |
| White | Any other white background |
| Mixed | White and Black Caribbean |
| Mixed | White and Black African |
| Mixed | White and Asian |
| Mixed | Any other mixed background |
| Asian or Asian British | Indian |
| Asian or Asian British | Pakistani |
| Asian or Asian British | Bangladeshi |
| Asian or Asian British | Any other Asian background |
| Black or Black British | Caribbean |
| Black or Black British | African |
| Black or Black British | Any other Black background |
| Chinese |  |
| Other ethnic group |  |
| Do not know |  |
| Prefer not to answer |  |

**Table S9:** Frequency of country of birth by cluster for Figure 4a. Proportion refers to the proportion within the cluster. Categories with proportion below 0:05 are not listed.

| Cluster | COB | Count | Proportion | Cluster | COB | Count | Proportion |
| --- | --- | --- | --- | --- | --- | --- | --- |
| n/a | England | <20 | 0.51 | 12 | Peru | 35 | 0.29 |
| n/a | Morocco | <20 | <0.1 | 12 | Ecuador | 24 | 0.20 |
| n/a | Sudan | <20 | <0.1 | 12 | Mexico | 21 | 0.17 |
| n/a | Libya | <20 | <0.1 | 12 | Bolivia | <20 | <0.2 |
| n/a | Wales | <20 | <0.1 | 12 | Colombia | <20 | <0.2 |
| 0 | Japan | 242 | 0.84 | 13 | Hong Kong | 459 | 0.22 |
| 0 | South Korea | 26 | 0.09 | 13 | China | 373 | 0.18 |
| 1 | Italy | 35 | 0.83 | 13 | Philippines | 321 | 0.16 |
| 1 | England | <20 | <0.2 | 13 | Malaysia | 314 | 0.15 |
| 2 | Finland | 136 | 0.92 | 14 | England | 194 | 0.49 |
| 3 | England | 1707 | 0.82 | 14 | Myanmar (Burma) | 24 | 0.06 |
| 4 | England | 2418 | 0.76 | 14 | Hong Kong | 23 | 0.06 |
| 4 | Scotland | 181 | 0.06 | 15 | Nepal | 123 | 0.80 |
| 4 | USA | 170 | 0.05 | 15 | Prefer not to answer | <20 | <0.1 |
| 5 | Iran | 502 | 0.31 | 16 | Spain | 330 | 0.39 |
| 5 | Iraq | 303 | 0.19 | 16 | Portugal | 282 | 0.33 |
| 5 | England | 169 | 0.10 | 16 | England | 56 | 0.07 |
| 5 | Cyprus | 163 | 0.10 | 17 | England | 355844 | 0.82 |
| 5 | Turkey | 135 | 0.08 | 17 | Scotland | 37490 | 0.09 |
| 6 | Egypt | 72 | 0.22 | 18 | Zimbabwe | 258 | 0.26 |
| 6 | Algeria | 70 | 0.21 | 18 | Congo | 144 | 0.14 |
| 6 | Morocco | 66 | 0.20 | 18 | Uganda | 126 | 0.13 |
| 6 | Libya | 37 | 0.11 | 18 | Kenya | 111 | 0.11 |
| 7 | India | 3019 | 0.33 | 18 | Zambia | 56 | 0.06 |
| 7 | Pakistan | 1344 | 0.15 | 18 | South Africa | 53 | 0.05 |
| 7 | Kenya | 1067 | 0.12 | 19 | Caribbean | 2268 | 0.31 |
| 7 | England | 743 | 0.08 | 19 | England | 2077 | 0.28 |
| 7 | Sri Lanka | 644 | 0.07 | 19 | Nigeria | 1017 | 0.14 |
| 8 | India | 140 | 0.33 | 19 | Ghana | 867 | 0.12 |
| 8 | Kenya | 124 | 0.29 | 20 | England | 3528 | 0.87 |
| 8 | Uganda | 81 | 0.19 | 21 | England | 8338 | 0.54 |
| 8 | England | 31 | 0.07 | 21 | Germany | 970 | 0.06 |
| 8 | Tanzania | 24 | 0.06 | 21 | Scotland | 938 | 0.06 |
| 9 | England | 553 | 0.60 | 22 | England | 697 | 0.66 |
| 9 | India | 190 | 0.21 | 22 | Caribbean | 79 | 0.08 |
| 10 | Somalia | 76 | 0.84 | 23 | Ethiopia | 57 | 0.33 |
| 10 | Prefer not to answer | <20 | <0.1 | 23 | Sudan | 50 | 0.29 |
| 11 | England | 50 | 0.58 | 23 | Eritrea | 44 | 0.25 |
| 11 | Wales | <20 | <0.2 | 24 | England | 3178 | 0.62 |
| 11 | France | <20 | <0.1 | 25 | England | 74 | 0.21 |
| 11 | Egypt | <20 | <0.1 | 25 | South Africa | 69 | 0.20 |
|  |  |  |  | 25 | Mauritius | 63 | 0.18 |
|  |  |  |  | 25 | Caribbean | 36 | 0.10 |

**Table S10:** Frequency of selected EB by cluster for Figure 4a. Proportions refer to the proportion within the cluster. Categories with proportions below 005 are not listed.

| Cluster | EB | Count | Proportion | Cluster | EB | Count | Proportion |
| --- | --- | --- | --- | --- | --- | --- | --- |
| n/a | Mixed, White and Black African | <20 | <0.3 | 12 | Other ethnic group, Other ethnic group | 90 | 0.74 |
| n/a | Mixed, Any other mixed background | <20 | <0.2 | 12 | Mixed, Any other mixed background | <20 | <0.2 |
| n/a | Other ethnic group, Other ethnic group | <20 | <0.2 | 12 | White, Any other white background | <20 | <0.2 |
| n/a | White, British | <20 | <0.2 | 13 | Chinese, Chinese | 1454 | 0.70 |
| n/a | White, Any other white background | <20 | <0.1 | 13 | Other ethnic group, Other ethnic group | 323 | 0.16 |
| n/a | Black or Black British, African | <20 | <0.1 | 13 | Asian or Asian British, Any other Asian background | 232 | 0.11 |
| 0 | Other ethnic group, Other ethnic group | 220 | 0.76 | 14 | Mixed, Any other mixed background | 114 | 0.29 |
| 0 | Asian or Asian British, Any other Asian background | 54 | 0.19 | 14 | Mixed, White and Asian | 93 | 0.23 |
| 1 | White, Any other white background | 39 | 0.93 | 14 | Other ethnic group, Other ethnic group | 61 | 0.15 |
| 2 | White, Any other white background | 145 | 0.98 | 14 | Asian or Asian British, Any other Asian background | 44 | 0.11 |
| 3 | White, British | 1585 | 0.76 | 14 | Chinese, Chinese | 31 | 0.08 |
| 3 | White, Any other white background | 407 | 0.20 | 14 | White, British | 31 | 0.08 |
| 4 | White, British | 1880 | 0.59 | 15 | Asian or Asian British, Any other Asian background | 63 | 0.41 |
| 4 | White, Any other white background | 993 | 0.31 | 15 | Other ethnic group, Other ethnic group | 63 | 0.41 |
| 4 | Other ethnic group, Other ethnic group | 239 | 0.08 | 15 | Not Available | 22 | 0.14 |
| 5 | Other ethnic group, Other ethnic group | 751 | 0.46 | 16 | White, Any other white background | 753 | 0.89 |
| 5 | White, Any other white background | 435 | 0.27 | 16 | White, British | 53 | 0.06 |
| 5 | Asian or Asian British, Any other Asian background | 229 | 0.14 | 17 | White, British | 412206 | 0.95 |
| 5 | White, British | 103 | 0.06 | 18 | Black or Black British, African | 773 | 0.77 |
| 6 | Other ethnic group, Other ethnic group | 223 | 0.68 | 18 | Other ethnic group, Other ethnic group | 190 | 0.19 |
| 6 | White, Any other white background | 47 | 0.14 | 19 | Black or Black British, Caribbean | 4143 | 0.56 |
| 6 | Mixed, White and Black African | <20 | <0.1 | 19 | Black or Black British, African | 2225 | 0.30 |
| 7 | Asian or Asian British, Indian | 5177 | 0.57 | 19 | Other ethnic group, Other ethnic group | 602 | 0.08 |
| 7 | Asian or Asian British, Pakistani | 1726 | 0.19 | 20 | White, British | 3778 | 0.93 |
| 7 | Asian or Asian British, Any other Asian background | 1049 | 0.12 | 20 | White, Any other white background | 221 | 0.05 |
| 7 | Other ethnic group, Other ethnic group | 498 | 0.06 | 21 | White, British | 7848 | 0.51 |
| 8 | Asian or Asian British, Indian | 419 | 0.99 | 21 | White, Any other white background | 7020 | 0.45 |
| 9 | Mixed, White and Asian | 432 | 0.47 | 22 | Mixed, White and Black Caribbean | 408 | 0.39 |
| 9 | White, British | 141 | 0.15 | 22 | Mixed, White and Black African | 254 | 0.24 |
| 9 | Asian or Asian British, Indian | 84 | 0.09 | 22 | Mixed, Any other mixed background | 111 | 0.11 |
| 9 | Mixed, Any other mixed background | 76 | 0.08 | 22 | Black or Black British, Caribbean | 106 | 0.10 |
| 9 | Other ethnic group, Other ethnic group | 53 | 0.06 | 22 | Other ethnic group, Other ethnic group | 69 | 0.07 |
| 10 | Black or Black British, African | 67 | 0.74 | 23 | Black or Black British, African | 95 | 0.54 |
| 10 | Other ethnic group, Other ethnic group | 22 | 0.24 | 23 | Other ethnic group, Other ethnic group | 64 | 0.37 |
| 11 | Mixed, Any other mixed background | 30 | 0.35 | 24 | White, British | 3416 | 0.67 |
| 11 | White, British | 20 | 0.23 | 24 | White, Any other white background | 646 | 0.13 |
| 11 | White, Any other white background | <20 | <0.2 | 24 | Other ethnic group, Other ethnic group | 285 | 0.06 |
| 11 | Mixed, White and Asian | <20 | <0.2 | 25 | Other ethnic group, Other ethnic group | 100 | 0.29 |
| 11 | Other ethnic group, Other ethnic group | <20 | <0.2 | 25 | Mixed, Any other mixed background | 83 | 0.24 |
|  |  |  |  | 25 | Mixed, White and Black African | 24 | 0.07 |
|  |  |  |  | 25 | Black or Black British, Caribbean | 23 | 0.07 |

**Table S11:** Comparing two phenotype models split by EB. One model (PCA) uses the top 40 PCs to estimate phenotypes, while the other (CLS) uses a cluster-smoothed phenotype estimate from Algorithm 1 in addition to the top 40 PCs. Values in the table are the average mean squared error with the average number of testing samples in a five-fold cross validation.

| phenotype | model | Any other mixed background | Any other white background | British | Chinese | Irish | Other ethnic group | Pakistani | White and Asian |
| --- | --- | --- | --- | --- | --- | --- | --- | --- | --- |
| Basophil count | CLS | 0.874 (n=194) | 0.835 (n=3058) | 1 (n=83497) | 0.511 (n=291) | 1.148 (n=2471) | 1.101 (n=837) | 1.126 (n=337) | 0.726 (n=156) |
| Basophil count | PCA | 0.942 (n=194) | 0.836 (n=3058) | 0.999 (n=83497) | 0.524 (n=291) | 1.151 (n=2471) | 1.114 (n=837) | 1.163 (n=337) | 0.763 (n=156) |
| BMI | CLS | 1.183 (n=198) | 1.051 (n=3150) | 0.984 (n=85947) | 0.649 (n=299) | 0.959 (n=2541) | 1.096 (n=858) | 0.936 (n=344) | 1.056 (n=160) |
| BMI | PCA | 1.219 (n=198) | 1.044 (n=3150) | 0.981 (n=85947) | 0.657 (n=299) | 0.961 (n=2541) | 0.991 (n=858) | 0.946 (n=344) | 1.159 (n=160) |
| Eosinophil count | CLS | 1.3 (n=194) | 0.917 (n=3058) | 0.974 (n=83497) | 1.171 (n=291) | 1.041 (n=2471) | 1.292 (n=837) | 1.661 (n=337) | 1.354 (n=156) |
| Eosinophil count | PCA | 1.384 (n=194) | 0.918 (n=3058) | 0.973 (n=83497) | 1.207 (n=291) | 1.044 (n=2471) | 1.268 (n=837) | 1.721 (n=337) | 1.451 (n=156) |
| Erythrocyte count | CLS | 1.092 (n=195) | 0.993 (n=3063) | 0.965 (n=83645) | 1.441 (n=291) | 0.989 (n=2477) | 1.251 (n=839) | 1.366 (n=337) | 1.069 (n=156) |
| Erythrocyte count | PCA | 1.157 (n=195) | 0.99 (n=3063) | 0.961 (n=83645) | 1.449 (n=291) | 0.992 (n=2477) | 1.23 (n=839) | 1.409 (n=337) | 1.147 (n=156) |
| FEV1 | CLS | 0.863 (n=181) | 0.867 (n=2896) | 0.918 (n=78597) | 0.861 (n=280) | 0.978 (n=2308) | 1.18 (n=786) | 1.264 (n=311) | 0.903 (n=148) |
| FEV1 | PCA | 0.82 (n=181) | 0.844 (n=2896) | 0.913 (n=78597) | 0.886 (n=280) | 0.98 (n=2308) | 0.961 (n=786) | 1.275 (n=311) | 0.882 (n=148) |
| FVC | CLS | 0.924 (n=181) | 0.852 (n=2896) | 0.899 (n=78597) | 0.937 (n=280) | 0.948 (n=2308) | 1.288 (n=786) | 1.39 (n=311) | 0.886 (n=148) |
| FVC | PCA | 0.866 (n=181) | 0.82 (n=2896) | 0.893 (n=78597) | 0.957 (n=280) | 0.951 (n=2308) | 1.074 (n=786) | 1.406 (n=311) | 0.861 (n=148) |
| Leukocyte count | CLS | 1.012 (n=195) | 0.977 (n=3063) | 0.987 (n=83644) | 0.939 (n=291) | 1.021 (n=2477) | 1.134 (n=839) | 0.861 (n=337) | 0.971 (n=156) |
| Leukocyte count | PCA | 1.067 (n=195) | 0.973 (n=3063) | 0.985 (n=83644) | 0.982 (n=291) | 1.026 (n=2477) | 1.06 (n=839) | 0.888 (n=337) | 1.024 (n=156) |
| Lymphocyte count | CLS | 1.01 (n=194) | 0.946 (n=3058) | 0.99 (n=83497) | 0.841 (n=291) | 0.987 (n=2471) | 1.046 (n=837) | 0.925 (n=337) | 0.972 (n=156) |
| Lymphocyte count | PCA | 1.039 (n=194) | 0.944 (n=3058) | 0.988 (n=83497) | 0.889 (n=291) | 0.99 (n=2471) | 1.025 (n=837) | 0.952 (n=337) | 1.032 (n=156) |
| Monocyte count | CLS | 1.508 (n=194) | 0.935 (n=3058) | 0.99 (n=83497) | 0.733 (n=291) | 1.071 (n=2471) | 1.027 (n=837) | 1.079 (n=337) | 1.178 (n=156) |
| Monocyte count | PCA | 1.632 (n=194) | 0.936 (n=3058) | 0.989 (n=83497) | 0.757 (n=291) | 1.076 (n=2471) | 1.007 (n=837) | 1.13 (n=337) | 1.215 (n=156) |
| Neutrophil count | CLS | 1.127 (n=194) | 0.966 (n=3058) | 0.977 (n=83497) | 0.974 (n=291) | 1.03 (n=2471) | 1.192 (n=837) | 0.886 (n=337) | 0.941 (n=156) |
| Neutrophil count | PCA | 1.174 (n=194) | 0.961 (n=3058) | 0.976 (n=83497) | 1.018 (n=291) | 1.033 (n=2471) | 1.09 (n=837) | 0.916 (n=337) | 0.994 (n=156) |
| Standing height | CLS | 1.027 (n=199) | 1.061 (n=3154) | 0.967 (n=86037) | 0.799 (n=299) | 0.936 (n=2544) | 1.058 (n=859) | 0.888 (n=344) | 0.873 (n=160) |
| Standing height | PCA | 0.993 (n=199) | 0.96 (n=3154) | 0.953 (n=86037) | 0.815 (n=299) | 0.932 (n=2544) | 0.914 (n=859) | 0.905 (n=344) | 0.896 (n=160) |
| Weight | CLS | 1.181 (n=198) | 1.032 (n=3151) | 0.978 (n=85979) | 0.673 (n=299) | 0.97 (n=2543) | 1.208 (n=859) | 0.913 (n=344) | 1.039 (n=160) |
| Weight | PCA | 1.193 (n=198) | 1.019 (n=3151) | 0.976 (n=85979) | 0.686 (n=299) | 0.972 (n=2543) | 1.001 (n=859) | 0.921 (n=344) | 1.137 (n=160) |

**Table S12:** Comparing two phenotype models split by EB. One model (PCA) uses the top 40 PCs to estimate phenotypes, while the other (CLS) uses a cluster-smoothed phenotype estimate from Algorithm 1 in addition to the top 40 PCs. Values in the table are the average mean squared error with the average number of testing samples in a five-fold cross validation.

| phenotype | model | African | Any other Asian background | Bangladeshi | Caribbean | Indian | White | White and Black African | White and Black Caribbean |
| --- | --- | --- | --- | --- | --- | --- | --- | --- | --- |
| Basophil count | CLS | 1.107 (n=607) | 0.739 (n=335) | 0.529 (n=41) | 0.96 (n=813) | 0.922 (n=1094) | 1.351 (n=106) | 0.87 (n=78) | 1.701 (n=115) |
| Basophil count | PCA | 1.105 (n=607) | 0.753 (n=335) | 0.668 (n=41) | 0.971 (n=813) | 0.928 (n=1094) | 1.741 (n=106) | 1.172 (n=78) | 1.846 (n=115) |
| BMI | CLS | 1.052 (n=630) | 0.801 (n=344) | 2.398 (n=43) | 1.214 (n=849) | 0.878 (n=1112) | 1.285 (n=108) | 1.288 (n=80) | 1.169 (n=118) |
| BMI | PCA | 1.06 (n=630) | 0.811 (n=344) | 0.942 (n=43) | 1.22 (n=849) | 0.882 (n=1112) | 1.476 (n=108) | 1.545 (n=80) | 1.232 (n=118) |
| Eosinophil count | CLS | 1.365 (n=607) | 1.756 (n=335) | 1.936 (n=41) | 1.106 (n=813) | 1.61 (n=1094) | 0.94 (n=106) | 1.469 (n=78) | 0.981 (n=115) |
| Eosinophil count | PCA | 1.388 (n=607) | 1.792 (n=335) | 2.479 (n=41) | 1.116 (n=813) | 1.626 (n=1094) | 1.156 (n=106) | 1.556 (n=78) | 1.008 (n=115) |
| Erythrocyte count | CLS | 1.506 (n=608) | 1.214 (n=337) | 1.243 (n=42) | 1.523 (n=816) | 1.355 (n=1097) | 1.099 (n=106) | 1.361 (n=78) | 1.219 (n=115) |
| Erythrocyte count | PCA | 1.517 (n=608) | 1.23 (n=337) | 1.654 (n=42) | 1.546 (n=816) | 1.354 (n=1097) | 1.511 (n=106) | 1.744 (n=78) | 1.316 (n=115) |
| FEV1 | CLS | 1.212 (n=577) | 1.232 (n=315) |  | 0.964 (n=775) | 1.027 (n=1053) | 1.274 (n=96) | 0.843 (n=73) | 0.999 (n=109) |
| FEV1 | PCA | 1.176 (n=577) | 1.155 (n=315) |  | 0.968 (n=775) | 1.007 (n=1053) | 1.515 (n=96) | 0.989 (n=73) | 1.081 (n=109) |
| FVC | CLS | 1.416 (n=577) | 1.414 (n=315) |  | 1.138 (n=775) | 1.174 (n=1053) | 1.071 (n=96) | 0.855 (n=73) | 1.009 (n=109) |
| FVC | PCA | 1.384 (n=577) | 1.305 (n=315) |  | 1.147 (n=775) | 1.154 (n=1053) | 1.355 (n=96) | 0.975 (n=73) | 1.122 (n=109) |
| Leukocyte count | CLS | 0.912 (n=608) | 0.938 (n=337) | 0.86 (n=42) | 1.192 (n=816) | 0.854 (n=1097) | 1.034 (n=106) | 1.195 (n=78) | 1.045 (n=115) |
| Leukocyte count | PCA | 0.902 (n=608) | 0.939 (n=337) | 1.199 (n=42) | 1.19 (n=816) | 0.865 (n=1097) | 1.233 (n=106) | 1.654 (n=78) | 1.199 (n=115) |
| Lymphocyte count | CLS | 0.932 (n=607) | 1.09 (n=335) | 0.908 (n=41) | 1.082 (n=813) | 1.032 (n=1094) | 0.945 (n=106) | 1.079 (n=78) | 0.999 (n=115) |
| Lymphocyte count | PCA | 0.944 (n=607) | 1.084 (n=335) | 1.298 (n=41) | 1.094 (n=813) | 1.047 (n=1094) | 1.026 (n=106) | 1.251 (n=78) | 1.108 (n=115) |
| Monocyte count | CLS | 0.913 (n=607) | 1.178 (n=335) | 1.205 (n=41) | 1.011 (n=813) | 0.951 (n=1094) | 1.122 (n=106) | 0.927 (n=78) | 0.96 (n=115) |
| Monocyte count | PCA | 0.916 (n=607) | 1.195 (n=335) | 1.512 (n=41) | 1.018 (n=813) | 0.963 (n=1094) | 1.383 (n=106) | 1.101 (n=78) | 1.099 (n=115) |
| Neutrophil count | CLS | 1.007 (n=607) | 0.945 (n=335) | 0.893 (n=41) | 1.3 (n=813) | 0.861 (n=1094) | 1 (n=106) | 1.172 (n=78) | 1.036 (n=115) |
| Neutrophil count | PCA | 0.995 (n=607) | 0.962 (n=335) | 1.284 (n=41) | 1.298 (n=813) | 0.868 (n=1094) | 1.202 (n=106) | 1.591 (n=78) | 1.178 (n=115) |
| Standing height | CLS | 0.975 (n=631) | 0.937 (n=345) | 0.772 (n=44) | 0.955 (n=850) | 0.959 (n=1114) | 1.05 (n=108) | 1 (n=80) | 1.072 (n=119) |
| Standing height | PCA | 0.978 (n=631) | 0.926 (n=345) | 1.08 (n=44) | 0.96 (n=850) | 0.931 (n=1114) | 1.119 (n=108) | 1.097 (n=80) | 1.109 (n=119) |
| Weight | CLS | 1.15 (n=630) | 0.874 (n=347) | 0.799 (n=43) | 1.23 (n=851) | 0.919 (n=1135) | 1.261 (n=108) | 1.26 (n=80) | 1.142 (n=119) |
| Weight | PCA | 1.15 (n=630) | 0.859 (n=347) | 0.956 (n=43) | 1.237 (n=851) | 0.911 (n=1135) | 1.411 (n=108) | 1.527 (n=80) | 1.185 (n=119) |

